# Targeting LxCxE cleft pocket of retinoblastoma protein in M2 macrophages inhibits ovarian cancer progression

**DOI:** 10.1101/2024.05.10.593562

**Authors:** Evgenii N. Tcyganov, Taekyoung Kwak, Xue Yang, Adi Narayana Reddy Poli, Colin Hart, Avishek Bhuniya, Joel Cassel, Andrew Kossenkov, Noam Auslander, Lily Lu, Paridhima Sharma, Maria De Grecia Cauti Mendoza, Dmitry Zhigarev, Mark Gregory Cadungog, Stephanie Jean, Sudeshna Chatterjee-Paer, David Weiner, Laxminarasimha Donthireddy, Bryan Bristow, Rugang Zhang, Vladimir A. Tyurin, Yulia Y. Tyurina, Hülya Bayir, Valerian E. Kagan, Joseph M. Salvino, Luis J. Montaner

## Abstract

Ovarian cancer remains a major health threat with limited treatment options available. It is characterized by immunosuppressive tumor microenvironment (TME) maintained by tumor- associated macrophages (TAMs) hindering anti-tumor responses and immunotherapy efficacy. Here we show that targeting retinoblastoma protein (Rb) by disruption of its LxCxE cleft pocket, causes cell death in TAMs by induction of ER stress, p53 and mitochondria-related cell death pathways. A reduction of pro-tumor Rb^high^ M2-type macrophages from TME in vivo enhanced T cell infiltration and inhibited cancer progression. We demonstrate an increased Rb expression in TAMs in women with ovarian cancer is associated with poorer prognosis. Ex vivo, we show analogous cell death induction by therapeutic Rb targeting in TAMs in post-surgery ascites from ovarian cancer patients. Overall, our data elucidates therapeutic targeting of the Rb LxCxE cleft pocket as a novel promising approach for ovarian cancer treatment through depletion of TAMs and re-shaping TME immune landscape.

**Statement of significance:** Currently, targeting immunosuppressive myeloid cells in ovarian cancer microenvironment is the first priority need to enable successful immunotherapy, but no effective solutions are clinically available. We show that targeting LxCxE cleft pocket of Retinoblastoma protein unexpectedly induces preferential cell death in M2 tumor-associated macrophages. Depletion of immunosuppressive M2 tumor-associated macrophages reshapes tumor microenvironment, enhances anti-tumor T cell responses, and inhibits ovarian cancer. Thus, we identify a novel paradoxical function of Retinoblastoma protein in regulating macrophage viability as well as a promising target to enhance immunotherapy efficacy in ovarian cancer.

## INTRODUCTION

Ovarian cancer remains a serious health problem worldwide and currently available treatment methods demonstrate limited efficacy (1). Therefore, it is of high priority to optimize immunotherapy approaches and by that to increase survival outcomes of cancer patients (2).

Ovarian cancer resistance to current immunotherapy treatments is in part mediated by the immunosuppressive tumor microenvironment (TME) and accumulation of tumor-associated macrophages (TAMs) blocking T cell infiltration to the tumor and inhibiting anti-tumor responses (3,4). Consequently, depletion or “re-programming” of TAMs aiming to enhance T cell functions is a major priority for effective cancer immunotherapy. However, the limited experimental approaches identified to deplete macrophages (clodronate, anti-CSF1R, etc.) have not advanced due to off-target effects and/or targeting systemic macrophage populations rather than preferentially targeting TAMs (5,6). Therapy approaches that could target TAMs/pro-tumor M2 subset while keeping anti-tumor M1 cells are expected to have a greater promise.

Retinoblastoma protein (Rb coded by *Rb1* gene) is a well-known tumor suppressor due to its role in cell cycle regulation. Rb function is mainly studied in tumor cells with high proliferation potential. Although studies also implicate Rb in the control of tumor cell death decisions (7,8), its role in the biology of non-tumor cell types such as non-proliferative mature myeloid cells in the TME still requires better characterization (8,9). For example, *Rb1* gene is highly expressed in macrophages (The Human Protein Atlas data), but its role in this cell population remains unknown. Our prior data in human macrophages support a role for Rb in the regulation of their cell death (10–13). Rb contains at least two pockets in its structure (a major cell cycle regulation pocket with A and B domains; second cleft pocket in domain B including the LxCxE binding motif). Consequently, Rb pocket regions interact with numerous adaptor proteins able to selectively control cell cycle and survival events (14,15). The functional consequences of these interactions in immune cells at various differentiation status are still poorly understood and to be determined.

In this study, we investigated the anti-cancer effects of targeting TAMs with a small molecule compound binding Rb LxCxE cleft pocket or by myeloid conditional knockouts. We found that targeting the Rb LxCxE cleft pocket can induce cell death in M2 macrophages and reshape the immune cell landscape towards anti-tumor responses resulting in inhibition of ovarian cancer progression.

## RESULTS

### Targeted disruption of Rb interaction with partner proteins at the LxCxE cleft pocket

Based on our prior data linking the Retinoblastoma protein (Rb) in human macrophages to regulation of their cell death (10–13), the high expression of *Rb1* in myeloid cells over other immune cells (in Human Protein Atlas dataset, https://www.proteinatlas.org) (**Fig. 1A**) and the role of the LxCxE cleft pocket in cell death control (15,16), we hypothesized that the disruption of Rb binding partners at the LxCxE cleft pocket could induce cell death in macrophages. Specifically, LxCxE **(Supplementary Fig. S1A)** cleft pocket of Rb mediates interactions with numerous adaptor proteins able to mediate cell death outcomes such as E2F1, BAX, ASK1 and HDAC1 (16–21). Due to previous data showing that the 1,2,4-thiadiazolidine-3,5-dione (TDZD)- containing AP-3-84 compound (**Supplementary Fig. S1B**) can bind Rb and disrupt HPV-E7 binding to the Rb LxCxE cleft pocket (22), we focused our analysis on further characterizing this molecule’s activity on Rb adaptor proteins binding the LxCxE cleft pocket. To first reconfirm AP-3-84 binding to Rb, we developed AP-8-239 molecule that represents AP-3-84 linked to fluorescein isothiocyanate (FITC) fluorophore. AP-8-239 bound to Rb in a concentration-dependent manner and was displaced by AP-3-84 demonstrating direct reaction with Rb (**Supplementary Fig. S1C-F**). As Cys706 and Cys712 were shown to be critical residues for Rb adaptor protein binding to the LxCxE cleft pocket (23–25), we envisioned that AP-3-84 can engage the LxCxE cleft pocket first in order to subsequently bind the reactive TDZD headgroup to Rb Cys706 (residue lying under pocket). Using an Rb mutant version where all cysteines except Cys706 and Cys712 were replaced with alanine residues (C-A mutant Rb380-785), we could demonstrate that (a) AP-8-239 and AP-3-84 still bound mutant Rb, (**Supplementary Fig. S1G-I**) similarly to wild type Rb protein as described above, and (b) that AP-3-84 could disrupt Rb interactions with LxCxE binding proteins E2F1, HDAC1, BAX in a dose-dependent fashion (**Supplementary Fig. S1J-L**). Importantly, MDM2 or HDAC4 proteins binding outside the LxCxE cleft at the main Rb A/B domain pocket (**Supplementary Fig. S1J,K**) were not affected. Further confirming specificity, HDAC inhibitors could not disrupt Rb-HDAC1 interaction (**Supplementary Table S1**). Taken together, we show AP-3-84 can selectively interfere with Rb adaptor proteins binding the LxCxE cleft pocket.

**Fig. 1.**
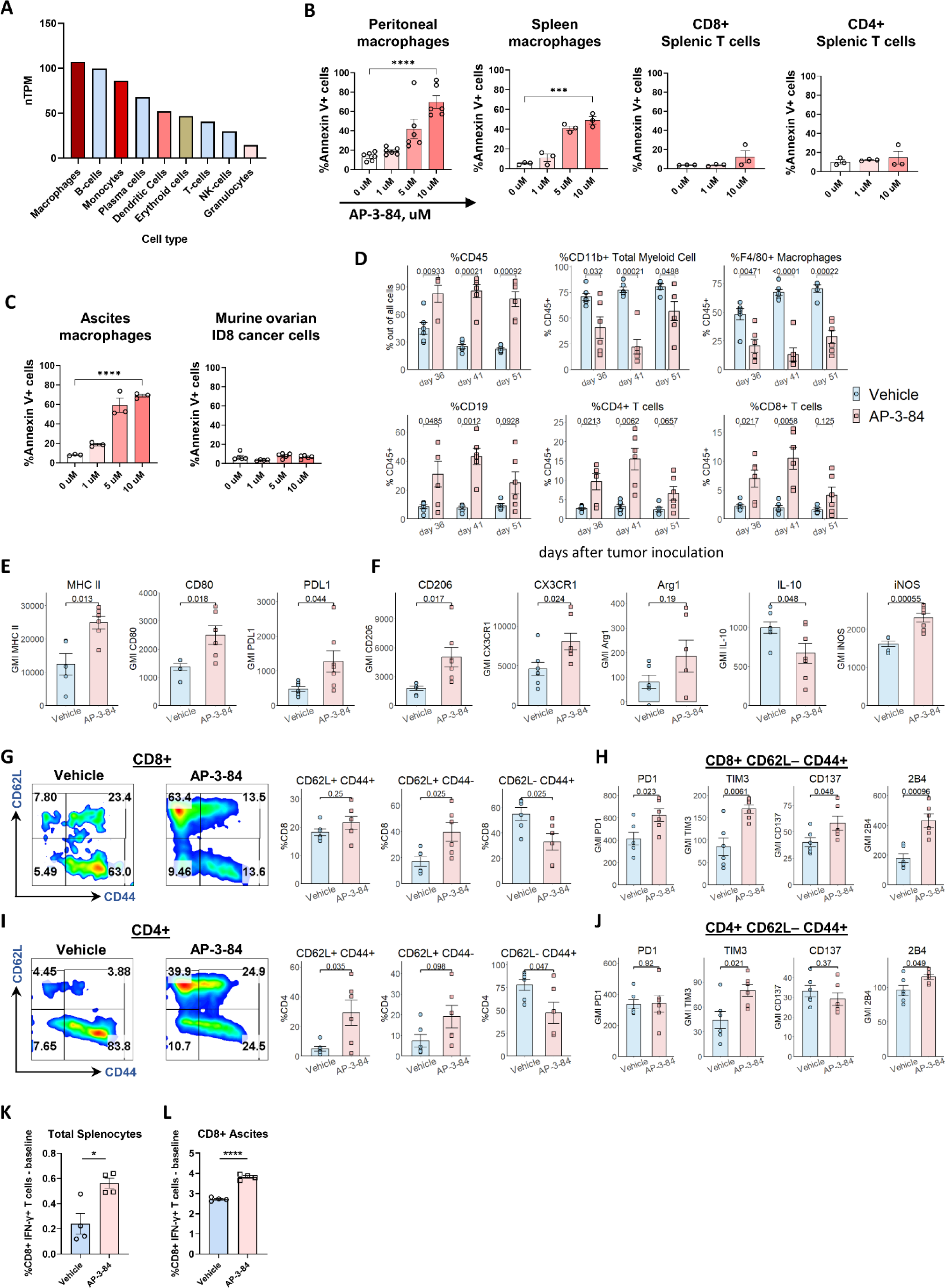
Therapeutic Rb targeting preferentially depletes macrophages and induces T cell infiltration in the ascitic tumor microenvironment of mice with ID8 ovarian cancer. **(A)** Expression of Rb1 gene in various immune cells. Data is obtained from The Human Protein Atlas dataset based on summarized meta-analysis of single-cell RNA sequencing of various human samples (https://www.proteinatlas.org/about/download, dataset #8). **(B)** Thioglycollate-induced peritoneal macrophages or spleen cells were incubated for 16-18 h with different doses of AP-3-84. The induction of cell death was measured in CD11b+ F4/80+ macrophages or CD3+CD4+ or CD3+CD8+ respective T cell subsets by Annexin V/Live-Dead staining using flow cytometry. n=3-6 per condition. **(C)** Ascites F4/80+ macrophages from ID8 ovarian cancer-bearing mice or original ID8 culture cells were incubated with AP-3-84 and analyzed for cell death induction as done in (**A).** n=3-5 per condition. **(D)** Immune cell subsets in ascites of AP-3-84-treated and control ID8-bearing mice at different stages of tumor progression analyzed by flow cytometry on the indicated days (also shown in **Supplementary Fig. S3B-E**). ID8 ovarian cancer model was performed as shown in **Supplementary Fig. S3A**. **(E,F)** AP-3-84 affects the expression of typical surface and intracellular macrophage markers on F4/80+ ascites macrophages. n=5-7 per group. **(G-J)** Ascites CD8+ (**G,H**) and CD4+ (**I,J**) T cells were analyzed for their differentiation state by CD62L/CD44 markers (**G,I**) and the expression of some co-inhibitory/co-stimulatory molecules on the effector CD62L–CD44+ subset was measured (**H,J**). n=5-6 per group. **(K,L)** The effects of AP-3-84 treatment on T cell tumor-specific responses in ID8-OVA-Luc-bearing mice. The frequencies of SIINFEKL-specific T cells in total splenocytes (**K**) and magnetically sorted ascites CD8+ T cells (**L**) were measured by intracellular flow cytometry. n=4 per group. Means +/– SEM are shown. Comparisons were conducted using the unpaired Student t-test; significance is indicated as *p<0.05, **p<0.01, ***p< 0.001, **** p<0.0001, ns – non- significant.

### Targeting the Rb LxCxE cleft pocket preferentially induces functional changes in macrophage cell death

Next, we tested the effects of Rb LxCxE cleft targeting on myeloid cell viability. We found that disruption of Rb by AP-3-84 induced cell death in thioglycolate-elicited peritoneal macrophages (**Fig. 1B** and **Supplementary Fig. S2A**) and in macrophages isolated *ex vivo* from the ascites of ID8 tumor-bearing mice (**Fig. 1C**). We observed the same effect in spleen macrophages, but not in T cell subsets (**Fig. 1B**). When we tested ID8 tumor cells, we did not reveal any significant effect of AP-3-84 on their viability (**Fig. 1C**). We also tested disruption of Rb by AP-3-84 in other commonly used ovarian cancer cell lines and did not detect any substantial effects on tumor cell viability (**Supplementary Fig. S2B**) supporting a refractory state in these cancer lines to disruption of Rb. Interestingly, we checked several different mesenchymal tumor cell lines and found similar refractoriness to Rb disruption of the LxCxE cleft pocket by AP-3-84 in contrast to TAMs (examples are shown in **Supplementary Fig. S2C**).

Next, we investigated whether Rb disruption of the LxCxE cleft pocket could impact cell division in ID8 cells as a proliferating tumor cell line in contrast to TAMs. Consistent with no effects on cell death, we found that tumor cell proliferation was not affected (**Supplementary Fig. S2D**). To further prove that effects associated with disruption of Rb are independent of upstream CDK4/6 signaling, we tested CDK4/6 palbociclib inhibitor and found that palbociclib did inhibit ID8 cell proliferation but had no effect on TAM viability (**Supplementary Fig. S2 E,F**). Thus, we interpret that disruption of Rb LxCxE cleft pocket can induce a preferential cell death in TAMs over ID8 cells or T cells without redundancy in activity with CDK4/6 or HDAC inhibitors.

### In vivo depletion of TAMs in ovarian cancer via Rb targeting re-shapes tumor microenvironment towards anti-tumor immune response

TAMs are associated with poor clinical outcomes in ovarian cancer (26,27). As Rb disruption of the LxCxE region demonstrated a capacity to induce cell death in ascites macrophages from ID8- bearing mice *in vitro*, we analyzed the *in vivo* effects of targeting Rb LxCxE cleft on immune cell composition in ovarian cancer TME. We injected mice with ID8 cells expressing luciferase (ID8-Luc), let the tumor to progress for 26 days, conducted a short course of seven daily AP-3- 84 treatments and isolated tumor ascites from AP-3-84-treated and control mice soon after the treatment end on day 36, then on day 41 and at progressed disease stage on day 51 (**Supplementary Fig. S3A**). Following treatment, we consistently observed a drop in tumor cell frequencies (CD45– cells) together with the accumulation of CD45+ leukocytes in the ascites with substantial subset changes in leukocyte composition (**Fig. 1D** and **Supplementary Fig. S3 B-E**). Specifically, we found a robust drop of total CD11b+ myeloid cells and F4/80+ macrophages, in particular (**Fig. 1D**), concurrently with a significant increase in B and T cells in the ascites of treated mice consistent with selective TAM depletion (**Fig. 1D** and **Supplementary Fig. S3C-E**).

We further analyzed how Rb LxCxE targeting changes the activation state of immune cells in TME. Macrophages remaining in TME after the 7-day treatment expressed higher levels of activation markers such as CD80, MHC II and PDL1 (**Fig. 1E**), but also retained high expression levels of such M2 markers as CD206 and CX3CR1 (**Fig. 1F**) consistent with an increase of M2 type TAMs in ovarian cancer (5,28), representing a “mixed” polarization phenotype. Surprisingly, we could document that AP-3-84 treatment induced a shift towards M1 type macrophages based on increased iNOS and PDL1 expression and reduced IL-10 production (**Fig. 1E,F**) suggesting a greater refractory state in M1 over M2 cells to Rb disruption as further tested below.

We also investigated the effects of 7-day treatment on T cell population changes (**Fig. 1D**). While increased in total frequency within ascites after treatment, both CD4+ and CD8+ T cell subsets had a lower degree of effector differentiation as reflected by the significant drop in CD62L–CD44+ effector cells and the accumulation of less differentiated CD62L+CD44- and CD62L+CD44+ subsets (**Fig. 1G,I**). Analysis of co-stimulatory/co-inhibitory T cell receptors further revealed that the CD8+CD62L-CD44+ effector subset in AP-3-84-treated animals had an increased level of co-inhibitory TIM3 and PD1 receptors together with simultaneous up- regulation of co-stimulatory CD137 (4-1BB) and 2B4 molecules (**Fig. 1H**). A similar trend was observed in the effector CD4+ counterpart subset (**Fig. 1J**). In this case, an increase in PD1 and TIM-3 expression may reflect the higher activation state of the effector CD8+ T cells, whereas CD137 and 2B4 may induce an enhanced secretory and proliferative activity of T cells.

To directly address the functional outcome of TAM depletion on anti-tumor specific T cell response, we used ID8 cells overexpressing chicken ovalbumin (ID8-OVA-Luc). In this model, we found an expansion of OVA-specific interferon-γ-producing T cells in splenocytes as well as ascitic CD8+ T cells 10 days after AP-3-84 treatment (**Fig. 1K,L**). Overall, our findings support that Rb LxCxE targeting in ovarian cancer induces a shift in the TME immune composition that can reduce the TAM immunosuppression barrier and allow for a greater anti-tumor T-cell response within the TME.

### Targeting Rb LxCxE cleft pocket preferentially affects M2 over M1-polarized macrophages

Based on the observed decrease of M2-like TAMs in the TME, we determined the Rb expression and effects of targeting Rb in M2 versus M1 macrophages as these subsets are described to play opposite roles in tumor progression (28,29). We first generated M0 (untreated), M1 or M2- polarized bone marrow-derived macrophages (BMDMs) as previously described (**Supplementary Fig. S3F**) (30). We analyzed gene expression by total RNA sequencing and generated differential M0/M1/M2 gene expression signatures (**Supplementary Table S2**). Using these signatures and principal component analysis (PCA), we characterized the polarization state of thioglycolate-induced macrophages and ascites macrophages from ID8 tumor-bearing animals. Interestingly, both thioglycolate-induced and ID8 ascites macrophages were most similar to M2 type cells whereas M0 and M1 type macrophages formed separate independent clusters regardless of AP-3-84 treatment (**Fig. 2A,B**). Supporting the similarity of TAMs and M2 BMDMs gene expression profiles, we also found that more than 85% TAMs in control ID8- bearing mice had an M2 type profile as judged by CD206 and CX3CR1 protein markers (**Fig. 2C**). Furthermore, *Rb1* gene expression was elevated in M2 BMDMs as compared to M1 (**Fig. 2D**). Flow cytometry analysis also confirmed the same difference at the protein level (**Supplementary Fig. S3G**). We co-incubated naïve peritoneal macrophages with the ascitic fluid obtained from ID8-bearing mice and again found the increase of the cleaved Rb expression (**Fig. 2E**). In addition, we found significantly higher level of Rb expression by immunofluorescence in the nucleus and cytoplasm of ascites macrophages when compared to ascites ID8 tumor cells (**Fig. 2F**), likely linked to the higher sensitivity of ascites macrophages to AP-3-84 (**Fig. 1C**). Consistent with the presence of cytoplasmic Rb expression in TAMs, we also found that Rb is mainly localized in the nucleus of M1-polarized BMDMs whereas it is largely expressed in the cytoplasm of M2 likely indicating Rb function outside the nucleus such as mediating interaction with mitochondrial proteins (19,31) (**Supplementary Fig. S3H**). Finally, we linked greater expression of Rb in M2-polarized BMDMs cells with a higher sensitivity to Rb targeting when compared to M1 (**Fig. 2G**). In sum, we interpret that (a) the higher RNA and protein expression of Rb and (b) the prevailing cytoplasmic distribution in M2-like TAMs contribute to the preferential M2-like cell targeting via LxCxE Rb modulation in TAMs within the ovarian cancer TME.

**Fig. 2.**
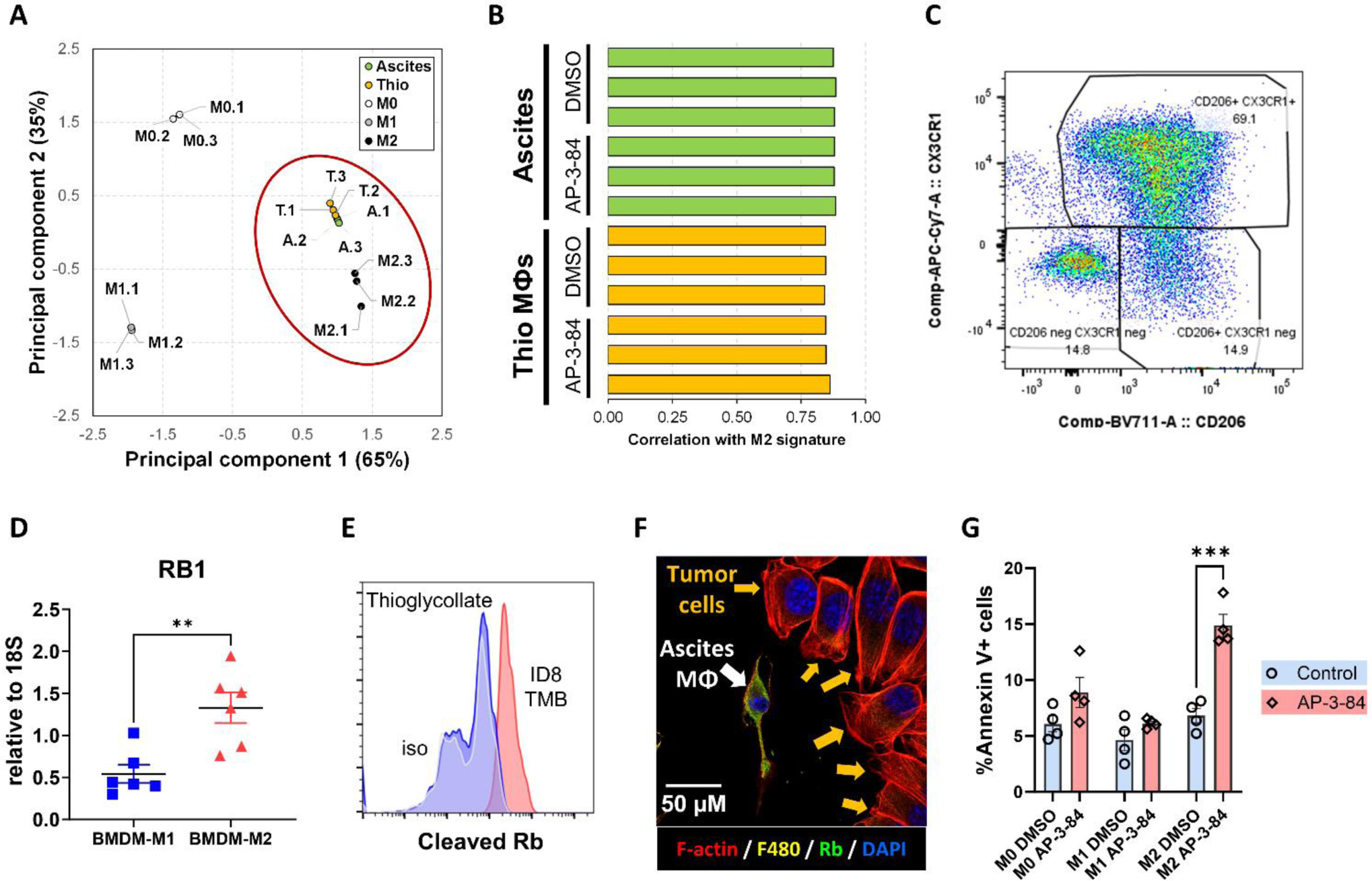
Therapeutic myeloid Rb targeting preferentially affects M2 type macrophages. (**A**) Principal component analysis comparing gene expression in thioglycolate-induced macrophages (“T”), ID8 ascites macrophages (“A”) and M0, M1 or M2-polarized bone marrow- derived macrophages. n=3 per group. (**B**) Correlation coefficients with M2 signature for thioglycolate-induced (“Thio МФs”) and ascites macrophages treated/non-treated with AP-3-84. n=3 per group. (**C**) Expression of CD206 and CX3CR1 M2 markers on ID8 ascites macrophages (isolated on day 36, gated on CD45+ CD11b+ F4/80+ Ly6G– cells). (**D**) qrtPCR analysis of *Rb1* gene expression in M1 or M2-polarized BMDMs. n=6 per group. (**E**) Control thioglycolate macrophages (blue) and macrophages pre-incubated with the ascitic fluid obtained from ID8-bearing mice (red) were analyzed for the cleaved Rb by intracellular flow cytometry staining. Representative experiment is shown. (**F**) The comparison of Rb expression (green) in the ascites macrophage with nearby ID8 tumor cells analyzed by confocal microscopy. Representative image is shown. (**G**) Generated BMDMs polarized to M0, M1 and M2 types were incubated overnight with 5 uM AP-3-84 and cell viability was assessed by Annexin V flow cytometry staining. n=4 per condition. Means +/–SEM are shown along with statistical significance for comparisons of corresponding groups by unpaired Student t-test, **p<0.01, ***p< 0.001.

### Targeting Rb LxCxE cleft pocket induces redundant cell death signaling pathways in macrophages

To determine what intracellular events are induced by myeloid Rb targeting, we analyzed gene expression changes in thioglycolate-elicited or ID8 ascites macrophages upon AP-3-84 treatment as well as in generated M0, M1 and M2 BMDMs. We found a major induction of a stress response program in both ascites and thioglycolate-elicited macrophages (**Fig. 3A,B,C** and **Supplementary Fig. S4A,B,C**). Stress response included unfolded protein response (UPR, also known as ER stress), NRF2-mediated oxidative stress response as well as apoptosis signaling induction. The induction of p53 target genes and inhibition of Rb1 pathway by AP-3-84 were readily detected (**Fig. 3B,C**).

**Fig. 3.**
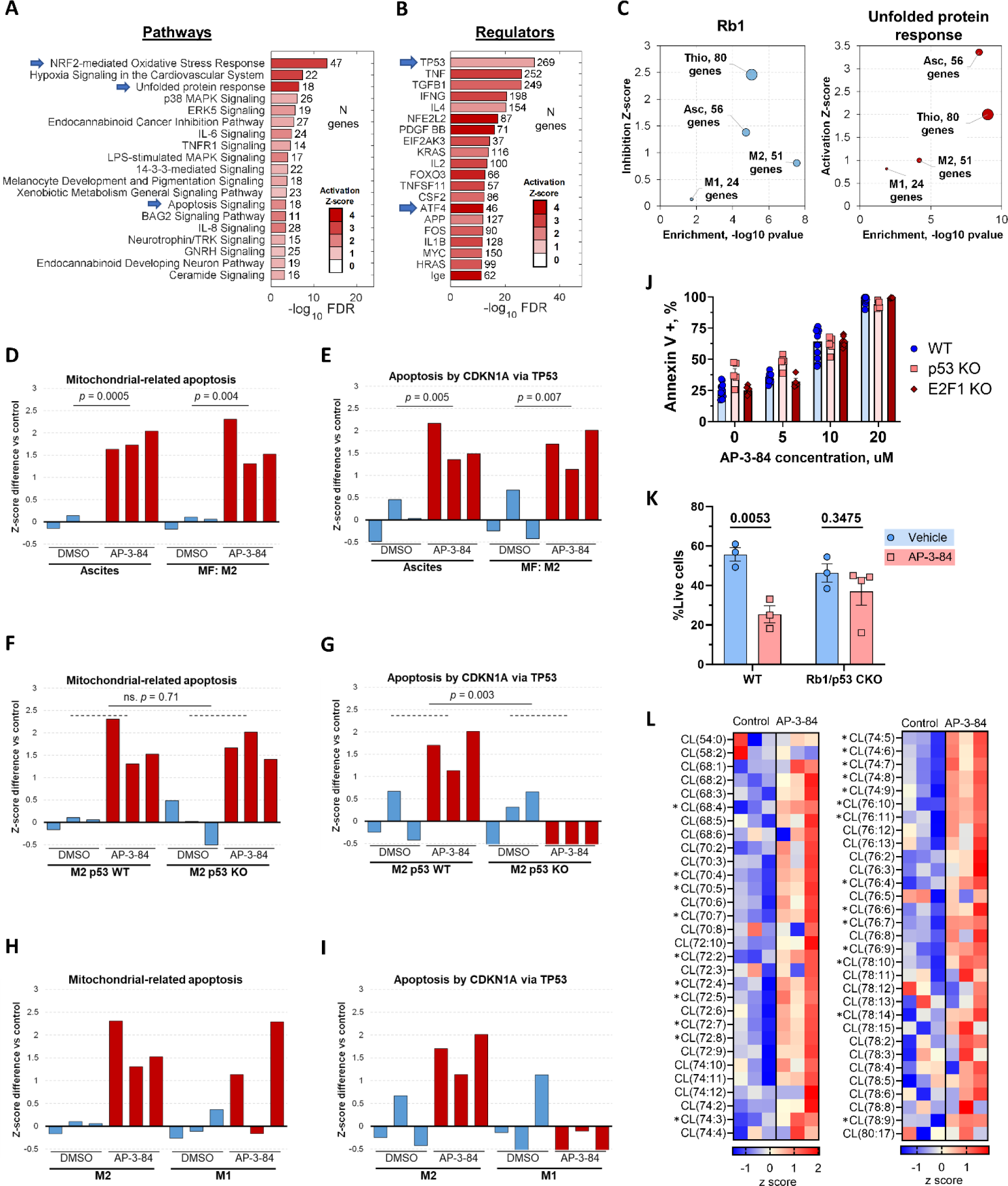
Myeloid Rb targeting initiates dramatic changes in macrophage gene expression profile with the induction of oxidative stress, unfolded protein response, and apoptosis signaling. ID8 ascites macrophages (isolated on day 42 after tumor inoculation), thioglycolate macrophages or M1 and M2-polarized BMDMs were treated *in vitro* with 10 uM AP-3-84 Rb modulator for 4 hours and their gene expression was analyzed by RNA sequencing (n=3 per condition). **(A, B)** IPA analysis of canonical pathways **(A)** and upstream gene regulators (**B**) most upregulated by AP-3-84 treatment in ID8 ascites macrophages. Characteristic hits are indicated by arrows. **(C)** Expression changes of target genes of RB1 and Unfolded protein response (UPR) signaling in different types of macrophages. Size of the bubbles is proportional to the number of target genes significantly affected by AP-3-84. **(D,E)** Single sample GSEA scores for induction of mitochondria-related apoptosis **(D)** or p53- related apoptosis **(E)** upon AP-3-84-treatment in ascites macrophages or M2-polarized BMDMs (shown as triplicates for each condition). **(F,G)** Similar to **D,E** GSEA score analysis in M2 wild-type versus p53 KO BMDMs. **(H,I)** Similar to **D,E** comparison for M1 and M2 wild-type BMDMs. **(J)** AP-3-84 induces similar cell death frequency in F4/80+ thioglycolate macrophages isolated from wild-type, p53 KO or E2F1 KO mice as measured by Annexin V flow cytometry staining. n=4-12 per condition. Means +/–SEM are shown. **(K)** Cre+ p53^floxed/floxed^ Rb1^floxed/floxed^ conditional knock-out (CKO) mice or their Cre negative wild-type (WT) littermates were injected with thioglycolate. After 4 days mice were treated with 5 mg/kg AP-3-84 intraperitoneally and peritoneal cells were collected in 3 hours with the following analysis of F4/80+ macrophage frequencies. n=3-4 per condition. **(L)** Cardiolipin species analysis by LC-MS/MS in thioglycolate-elicited wild-type F4/80+ macrophages upon AP-3-84 treatment. Heatmap demonstrating the differences in the contents of individual cardiolipin species is shown (n=3 per group). P-values for unpaired Student t-test comparisons are shown as exact values (D-I,K) or as *p<0.05 (L).

Using published datasets related to cell death (32), we found 2 main gene signatures induced by targeting Rb: mitochondria-related (GO:1901029) and p53/p21-mediated (33) cell death pathways. These pathways were clearly detected in ascites and M2 polarized macrophages upon treatment with AP-3-84 (**Fig. 3D,E**). To determine whether p53 controls both pathways, we generated M2 macrophages using bone marrow from p53 knock-out (p53 KO) mice and conducted similar analysis. Using PCA comparison of M0/M1/M2 BMDMs generated from wild type or p53 KO mice, we first established that p53 KO did not affect M1/M2 polarization in macrophages (**Supplementary Fig. S4D**) and that the p53-driven AP-3-84-induced gene expression was canceled in p53 KO M2 cells (**Fig. 3G**). Interestingly, the mitochondria-related cell death pathway was still detected (**Fig. 3F**) establishing the presence of a p53-independent pathway for AP-3-84-induced cell death signals. Finally, we compared these two signaling pathways in M2 versus M1 cells. In agreement with M1 resistance to AP-3-84 treatment, both signaling pathways were less pronounced in M1 cells upon AP-3-84 treatment (**Fig. 3H,I**). Consistent with the redundancy of cell death pathways induced by myeloid Rb targeting, p53 KO thioglycolate macrophages were still sensitive to AP-3-84 (**Fig. 3J**). As E2F1 is one of the main Rb partner proteins potentially involved in apoptosis regulation (16), we also tested AP-3-84 sensitivity of thioglycolate macrophages isolated from E2F1 KO mice. We found no difference with wild type cells (**Fig. 3J**).

Finally, to conclusively establish that Rb targeting induces cell death in peritoneal macrophages we took advantage of mice with conditional *Rb1/p53* genes knock-out in myeloid cells. For that we used LysM Cre+ Rb1^floxed/floxed^ p53^floxed/floxed^ mice (Rb1/p53 CKO). We utilized their Cre negative wild-type (WT) littermates as a control. We injected thioglycolate into those mice to induce peritoneal macrophage accumulation before in vivo exposure to AP-3-84 or vehicle control, collected peritoneal cells and analyzed macrophage frequencies. We found that single AP-3-84 treatment significantly reduced the presence of F4/80+ macrophages in the peritoneum of wild-type mice whereas peritoneal macrophages in Rb1/p53 CKO mice were not significantly affected (**Fig. 3K**). Having independently established that the p53 gene knock-out did not affect macrophage sensitivity to AP-3-84, we conclude that AP-3-84 effects are mediated via Rb targeting in the Rb1/p53 CKO.

### Targeting Rb LxCxE cleft pocket induces phospholipid changes associated with apoptosis and mitochondria-driven cell death outcomes in macrophages

Our previous work and other studies demonstrated phospholipids involvement in regulation of several cell death programs through peroxidation of polyunsaturated fatty acid (PUFA)- phospholipids (34,35). Cardiolipin (CL) is a characteristic active participant in apoptosis (36), while phosphatidylethanolamine (PE) and phosphatidylcholine (PC) contribute to ferroptosis and necroptosis (37), respectively. We performed global LC/MS-based quantitative phospholipidomics analysis of thioglycolate-induced and ascites macrophages to estimate AP-3- 84-mediated changes. We found that cardiolipins (CL) (**Fig. 3L and Supplementary Fig. S4E**) as well as phosphatidylcholines (PC), phosphatidylethanolamines (PE) and phosphatidylinositols (PIs) (**Supplementary Fig. S4I**) were upregulated by AP-3-84 in thioglycolate macrophages. AP-3-84 affected CL composition resulting in significant increase of highly oxidizable PUFA- CL species with 2-14 double bonds (**Fig. 3L**). This is not unexpected, given that AP-3-84 may target the Rb/E2F1 complex heavily involved in regulation of phospholipid metabolism, and PUFA-CL may undergo substantial oxidation leading to cell death (21,38). Similarly, PE, PC and PI species were sharply elevated mostly due to increased contents of PUFA-PE and PUFA-PC (**Supplementary Fig. S4I** and data not shown). AP-3-84 treatment of ID8 ascites macrophages also resulted in the increased CL and PE content (**Supplementary Fig. S4F-H**) with no changes in PC and PI (**Supplementary Fig. S4H**). Importantly, no AP-3-84-induced changes in non-mitochondrial phospholipid, phosphatidylserine, were detected in both types of macrophages, suggesting that mitochondria lipid changes were mainly involved in AP-3-84-induced response (**Supplementary Fig. S4H,I**). Overall, therapeutic Rb modulation strongly affected highly oxidizable PUFA-species of mitochondria-specific CL as well as PUFA-PC and PUFA-PE present in all cell compartments. This data further supports earlier gene expression data by documenting effects of myeloid Rb targeting of the LxCxE cleft pocket on cardiolipins and mitochondria-mediated cell death pathways.

We further confirmed gene expression findings on the effects of myeloid Rb targeting at the protein level. We could detect p53 induction and its target p21 in ascites macrophages shortly after *in vitro* AP-3-84 treatment (**Fig. 4A**). To verify p53 up-regulation by AP-3-84, we also applied immunofluorescence microscopy and detected consistent enhancement of p53 signal in AP-3-84-treated thioglycolate macrophages (**Supplementary Fig. S5A**). Hsp27 induction as a feature of ongoing ER stress was also readily detectable (**Fig. 4A**). In agreement with the initiation of NRF2-mediated oxidative stress response as noted in gene expression data, we could also detect the induction of reactive oxygen species (ROS) in thioglycolate macrophages as early as 2 hours upon the treatment (**Fig. 4B,C**).

**Fig. 4.**
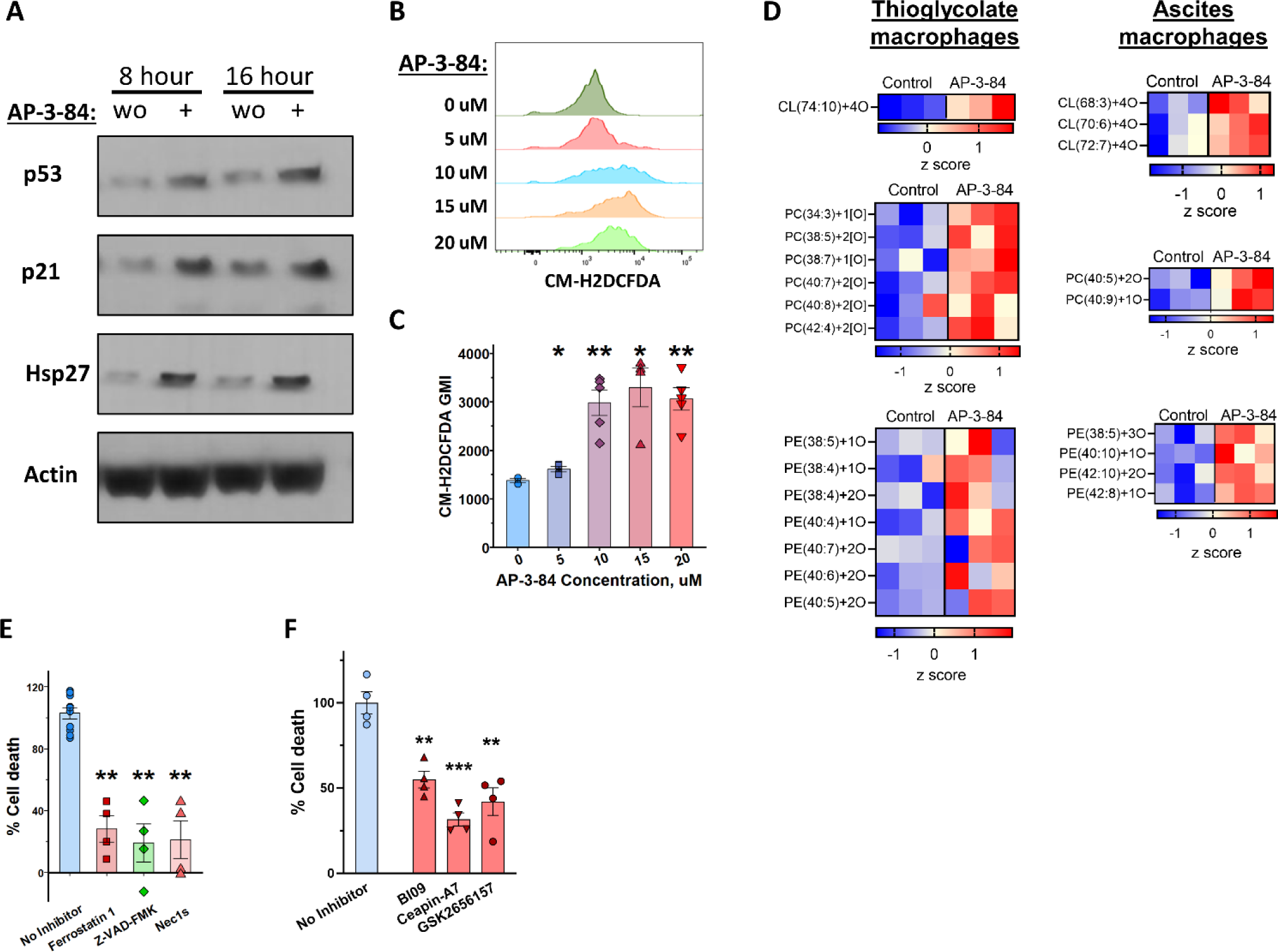
Rb modulation induces major stress response programs and regulates cell death pathways in macrophages. **(A)** Expression of apoptosis-related proteins measured by western blot in ascites macrophages treated *in vitro* with 10 µM AP-3-84 for the specified time. Representative data of at least two independent experiments is shown. **(B,C)** Induction of ROS in thioglycolate-induced macrophages after 2 h incubation with 10 uM AP-3-84 as measured by CM-H2DCFDA dye and flow cytometry Representative histograms (**B**) and summarized data of GMI fluorescence (**C**) are shown (n=4 per condition). **(D)** AP-3-84 treatment resulted in accumulation of oxygenated CL, PC and PE species in thioglycolate-induced (left) and ID8 ascites (right) macrophages as measured by LC-MS/MS (n=3 per condition). **(E,F)** Inhibition of AP-3-84-induced cell death by the inhibitors specific for general cell death programs **(E)** or for ER-stress pathway **(F)** in M2-polarized thioglycolate-induced macrophages, n=4-9 per condition. Means +/–SEM are shown. P-values for unpaired Student t-test comparisons with control cells are indicated: ns – no significant, *p<0.05, **p<0.01, ***p< 0.001, **** p<0.0001.

Initiation of different cell death pathways is accompanied by generation of specific lipid-derived signals. Apoptosis, ferroptosis and necroptosis are all associated with accumulation of oxygenated species of CL, PE, and PC, respectively (37). Using redox lipidomics approach, we found that Rb modulation by AP-3-84 resulted in accumulation of oxidized CL species and several oxygenated PE and PC species in both thioglycolate-elicited and ascites macrophages (**Fig. 4D**). This is consistent with the simultaneous involvement of three above-mentioned regulated cell death programs, apoptosis, ferroptosis and necroptosis, in the response of macrophages to therapeutic Rb LxCxE cleft pocket targeting.

To further characterize the role for Rb modulation in cell death induction, we incubated IL-4 treated M2-polarized thioglycolate macrophages with the inhibitors specific for different cell death pathways. We found a robust cancellation of cell death by the inhibitors of necroptosis, apoptosis and ferroptosis (Nec-1s, Z-Vad-FMK and Ferrostatin-1, respectively) (**Fig. 4E**) confirming the redundancy of cell death pathways induced by AP-3-84. As ER stress can initiate cell death (39) and was clearly induced by AP-3-84, we tested the inhibitors specific for three main ER stress pathways (BI09, Ceapin-A7 and GSK2656157 as respective IRE1α, ATF6 and PERK inhibitors) in M2-polarized thioglycolate macrophages. ER stress inhibition also significantly reversed AP-3-84-induced cell death in macrophages indicating its direct involvement (**Fig. 4F**).

We also tested inhibiting ASK1, known to bind to the LxCxE cleft pocket and play a role in apoptosis (40) and found no significant effect (data not shown). As other TDZD-derivative compounds (Tideglusib and TDZD-8) were previously shown to induce cell death in tumor cells through GSK3β inhibition (41,42), we also tested non-covalent GSK3β inhibitor Laduviglusib for induction of cell death in thioglycolate macrophages and found no effect on cell viability (data not shown). Taken together, we interpret that Rb disruption of the LxCxE cleft pocket induces redundant cell death programs in M2 macrophages with ER stress as one of the major initiating intracellular mechanisms leading to cell death.

### Rb targeting in myeloid cells inhibits ovarian cancer growth in a T cell-dependent manner

As targeting Rb reshaped ovarian cancer TME towards enhanced T cell responses, we assessed the effects of that on the tumor growth. As above, we conducted a short course of AP-3-84 treatment in the established ID8-Luc tumor model and monitored tumor growth by *In Vivo* Imaging System (IVIS) (**Supplementary Fig. S3A**). We found that AP-3-84 treatment significantly inhibited subsequent ID8 tumor growth (**Fig. 5A**) as well as the other tested models of ovarian cancer (BPPNM and KPCA, **Supplementary Fig. S6A,B**, respectively). In ID8 model, by IVIS we did note a temporal bioluminescence signal increase at day 35 (**Fig. 5A**), which we assigned to tumor lysis rather than a tumor growth as we detected a drop of tumor cell frequencies in the ascites of treated animals on the same day (day 35) and later time-points (**Supplementary Fig. S3B**). The effect of Rb targeting was also more pronounced when we used ID8-OVA cells expressing chicken ovalbumin, which is known to elicit robust specific T cell responses. A similar course of AP-3-84 treatment induced an earlier tumor burden reduction in ID8-OVA-bearing mice compared to the ID8 model likely due to the presence of a “stronger” OVA-tumor antigen (**Fig. 5B**) and consistent with enhanced anti-OVA T cell responses as shown above (**Fig. 1K,L**). As expected, we also found that tumor burden was positively correlated with the higher macrophage and total CD11b+ myeloid cell presence and negatively correlated with T cell infiltration (**Fig. 5C**) confirming that TME changes after AP-3-84 treatment are consistent with lower tumor burden.

**Fig. 5.**
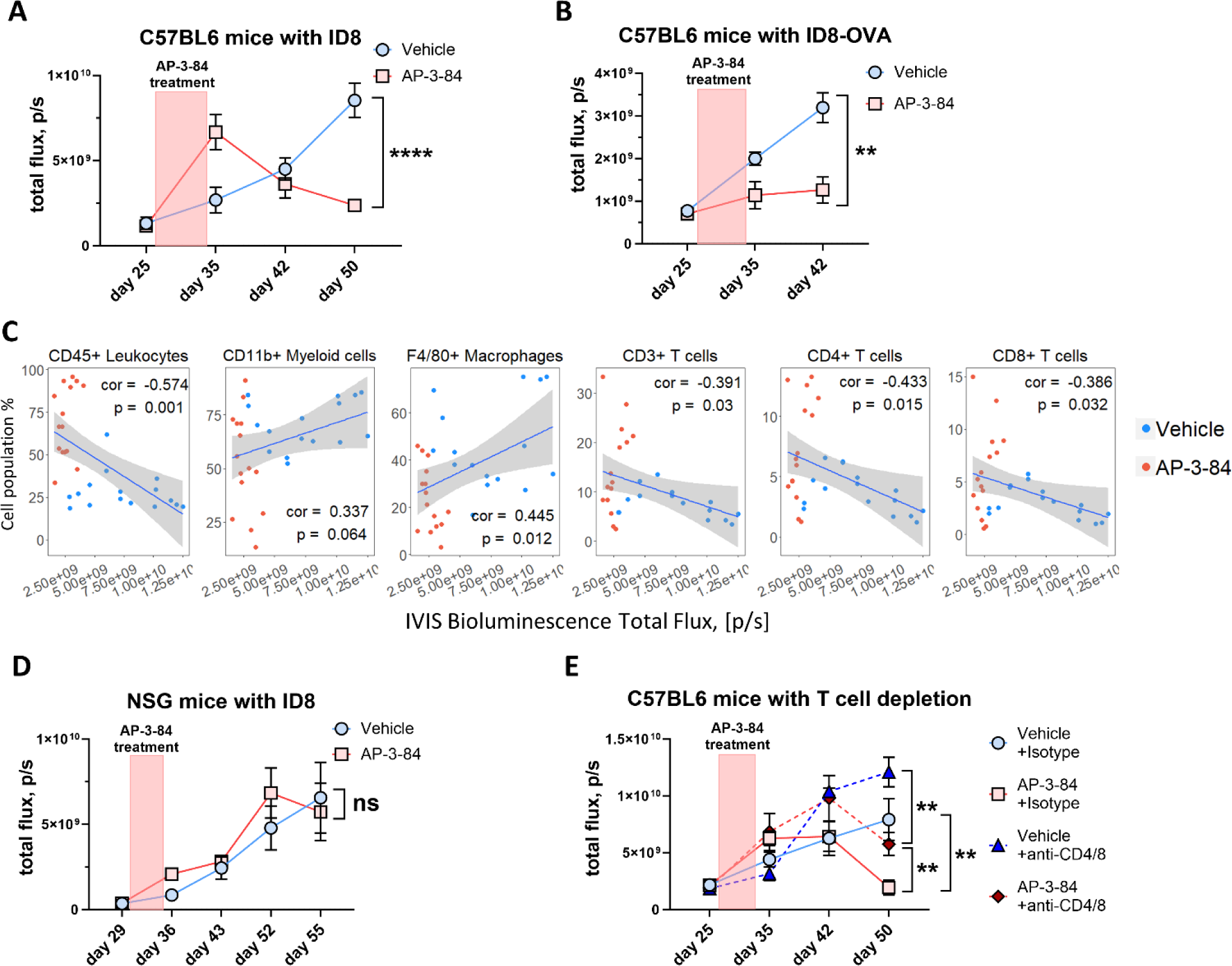
Therapeutic Rb targeting delays the ovarian cancer progression in immune- mediated manner. ID8 ovarian cancer model was performed as shown in **Supplementary Fig. S3A**. **(A)** AP-3-84 treatment delays ID8-Luc tumor progression. n=6-13 per time-point per group. **(B)** AP-3-84 treatment delays ID8-OVA-Luc tumor progression. Tumor burden was measured by IVIS as in **(A)**. n=5-7 per time-point per group. **(C)** Pearson correlation of IVIS measurements of tumor burden at pre-lethal stage (day 51) with the frequencies of the indicated immune cell subsets. The correlation coefficient (cor) and p- value are shown, n=31. **(D)** AP-3-84 treatment of ID8-bearing NSG mice conducted same as in (**A**). n=7 per condition. **(E)** AP-3-84 treatment of immunocompetent B6 mice combined with anti-CD4/CD8 T cell depletion (started simultaneously with AP-3-84 treatment on day 27). n=6-7 per group per time- point. Means +/– SEM are shown. Comparisons were conducted using the unpaired Student t-test; significance is indicated as *p<0.05, **p<0.01, ***p< 0.001, **** p<0.0001, ns – non- significant.

To directly establish a role for anti-tumor immune responses in decreasing tumor burden, we repeated the experiment with the same design as before (**Supplementary Fig. S3A**), but in NSG immunodeficient mice (those mice lack mature T cells, B cells and NK cells). In contrast to immunocompetent mice, we did not observe the tumor burden reduction in the treated NSG tumor-bearing animals (**Fig. 5D**). To further confirm the role of T-cell responses, we conducted anti-CD4/CD8 T cell depletion started simultaneously with AP-3-84 treatment. AP-3-84-treated mice that underwent antibody-mediated T-cell depletion had a significantly higher tumor burden when compared to intact AP-3-84 treated animals. However, we did observe that AP-3-84- treated mice with T cell depletion still had lower tumor burden than control animals that received anti-T cell depleting antibodies (**Fig. 5E**) indicative of T-cell-independent effects of TAM depletion by AP-3-84. Taken together, we interpret that Rb disruption of the LxCxE cleft pocket by AP-3-84 results in a substantial decrease of tumor burden via modulation of immune-mediated mechanism as a consequence of TAM depletion.

### Targeting the LxCxE pocket of Rb in the tumor microenvironment is not redundant to the clinically available CDK4/6 inhibitor palbociclib

To establish that *in vivo* effects of myeloid Rb targeting on the TME are not redundant to the other clinically available therapies decreasing tumor burden by targeting the main Rb pocket function in tumor cells with palbociclib CDK4/6 inhibitor, we tested AP-3-84, palbociclib or their combination in the ID8 model (**Supplementary Fig. S6C**). We observed similar effects by each single-agent therapy, albeit by different mechanisms, in reducing the tumor burden while their combination resulted in the strongest tumor growth inhibition (**Supplementary Fig. S6D**). As expected, AP-3-84 alone or palbociclib alone reduced macrophage frequency in the ascites presumably by different mechanisms: due to direct anti-TAM activity of AP-3-84 or due to the reduction of tumor burden by palbociclib. However, only AP-3-84 or its combination with palbociclib, induced tumor ascites infiltration with CD4+ and CD8+ T cells (**Supplementary Fig. S6E**). We interpret that disruption of Rb LxCxE cleft pocket by AP-3-84 can result in greater anti-tumor reduction *in vivo* via modulation of immune cell changes in TME not otherwise achieved by palbociclib direct anti-tumor cytotoxicity alone.

### Rb targeting depletes TAMs in ex vivo specimens from ovarian cancer patients

We investigated whether our TAM-depleting approach via myeloid Rb targeting could be relevant for ovarian cancer patients. First, we sought to determine whether patient TAMs express *Rb1* in the TME. Using the TCGA database for ovarian cancer patients and GTEx data for healthy donors, we found a stronger significant correlation between CD68 and *Rb1* in the ovarian tumor tissue (**Fig. 6A**) when compared against healthy ovarian tissue (**Fig. 6B**). We then compared *Rb1* expression in the tumor site and the adjacent healthy ovary tissue and further confirmed that *Rb1* is significantly higher expressed inside the tumor (**Fig. 6C**). We then used the CIBERSORT algorithm to predict M2 macrophage abundance in the tumor based on gene expression and then correlated that with *Rb1* expression in the same patient samples. Again, we found a strong correlation between M2 levels and *Rb1* supporting what we revealed in ID8 ascites TAMs expressing high Rb levels (**Fig. 6D**). Using an ovarian cancer patient cohort from the TCGA, we correlated *Rb1* gene expression levels with the predicted abundance of different immune cell subsets inside the tumor. We found that tumor M2 macrophages and neutrophils were predicted to have the highest levels of *Rb1* expression (**Fig. 6E**). Based on these data, we checked AP-3-84 effect on thioglycolate-induced murine neutrophil viability and found neutrophils to be significantly less sensitive to AP-3-84 than thioglycolate-induced macrophages (data not shown).

**Fig. 6.**
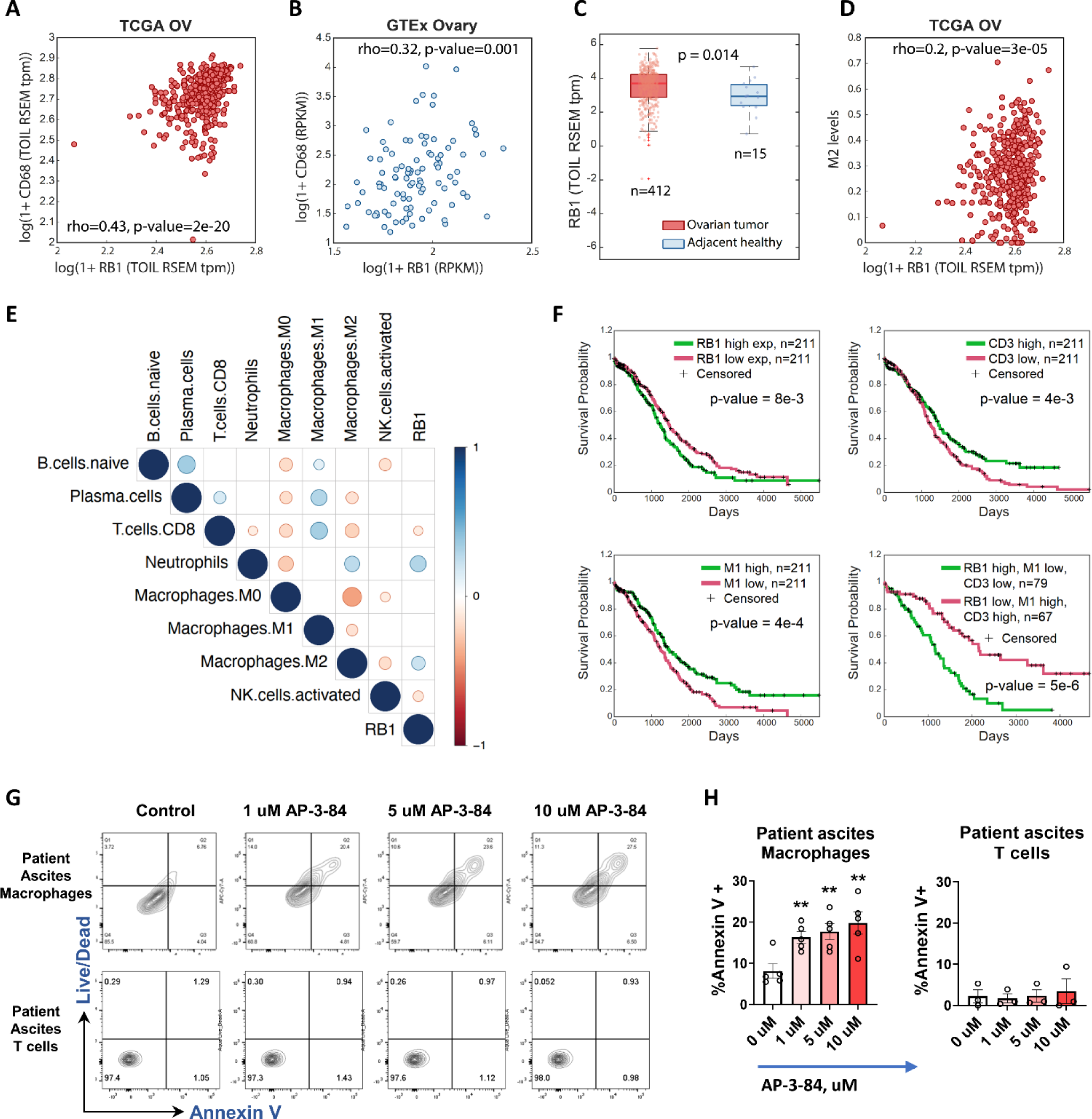
*Rb1* gene is expressed in TAMs of cancer patients and is associated with worse prognosis of ovarian cancer outcome. **(A,B)** Scatter plots correlating gene expression levels of *Rb1* gene with CD68 in TCGA ovarian cancer **(A)** and in GTEx healthy ovary **(B)**. The Spearman correlation coefficient and p-values are shown. (**C**) Comparison of *Rb1* gene expression levels in ovarian tumors (TCGA data) and adjacent healthy tissues. Medians with interquartile range and the rank-sum p-value are shown. (**D**) Correlation of *Rb1* gene expression to CIBERSORT-inferred M2-type macrophage frequencies in TCGA ovarian cancer dataset. The Spearman correlation coefficient and p-value are shown. (**E**) Heatmap of the rank correlation coefficients for *Rb1* expression and CIBERSORT-inferred immune cell abundances in TCGA ovarian cancer samples. (**F**) Kaplan-Meier survival curves comparing subgroups in TCGA ovarian cancer patients based on *Rb1* expression, or M1 and CD3+ T cell CIBERSORT-inferred abundances. The log-rank p-values are indicated. **(G,H)** AP-3-84 induces cell death in macrophages, but not in T cells in post-surgery ascites from ovarian cancer patients, n=5 per condition. Conditions were compared using unpaired Student t- test, **p<0.01.

Lastly, we analyzed the available cohort of ovarian cancer patients asking whether *Rb1* expression could be an outcome predictor. As hypothesized, we found that higher *Rb1* expression on its own was a predictor of poorer clinical outcome (**Fig. 6F**). However, the prediction was much stronger when high *Rb1* expression was combined with low M1 macrophage and low CD3+ T cell abundance (**Fig. 6F**).

As the Rb protein family also includes 2 more additional members, RBL1 and RBL2, which may have overlapping functions with Rb (14), we checked whether there might be analogous correlation of *RBL1* and *RBL2* with the survival outcome of ovarian cancer patients. In contrast to *Rb1* gene, we did not detect the same association for *RBL1* or *RBL2* genes, underlining *Rb1* as a major factor in this relation (**Supplementary Fig. S7A,B,C**).

Considering that Rb in TAMs is a potential predictor of poor clinical outcome in ovarian cancer patients, we decided to check whether our same therapeutic myeloid Rb targeting approach could be used for TAM depletion in the human TME. We obtained same-day post-surgery ascites from debulking surgeries of ovarian cancer patients and incubated ascites cells with different concentrations of AP-3-84. We observed that similarly to our murine ID8 model, AP-3-84 exposure is able to induce cell death in ascites macrophages from patients, with a much reduced to absent effect on T cells (**Fig. 6G,H**). Overall, this data supports prior animal model data indicating that therapeutic myeloid Rb targeting may represent an attractive immunotherapy strategy for TAM depletion in ovarian cancer patients.

## DISCUSSION

TAMs have been recognized as a factor inhibiting anti-tumor T cell functions and limiting immunotherapy efficacy in ovarian cancer (5). In this study, we identify a novel role for the LxCxE cleft pocket of Rb protein in determining TAM cell death outcomes in M2-like macrophages within the TME in either murine ID8 model of ovarian cancer or in ascites macrophages from ovarian cancer patients. Importantly, the depletion of M2-like TAMs via Rb disruption of the LxCxE cleft pocket resulted in a shift to M1 macrophages, enhanced T cell infiltration into TME and consequential inhibition of ID8 tumor growth. We establish a higher Rb expression in M2 or TAMs, when compared to M1 cells, which supports prior reports showing BCG-induced M1 type cells reducing their Rb expression (43) and M2 tumor-infiltrating macrophages with higher Rb expression (44). Consistent with M2 macrophages as “pro-tumor” cells inhibiting anti-tumor T cell responses in ovarian cancer patients (28,29), we found that higher abundance of M2-polarized cells expressing higher Rb levels is associated with a poorer clinical outcome.

Thus far, Rb protein has been primarily considered under its cell cycle-regulating activities rather than playing a central role in maintaining viability in differentiated non-dividing immune cells (8,45). Consistent with our data, it was previously shown that Rb inactivation can not only induce cell cycle, but also trigger the apoptosis pathway in p53-dependent or independent manner (8,16,20). Our study now demonstrates that disruption of the Rb LxCxE pocket can preferentially induce cell death within M2 macrophages while not having a similar activity in the other cell types such as tumor cells or T cells. Altogether, this supports a new role for Rb acting to maintain M2-like TAM myeloid cell survival in the TME.

Our data also confirms previously reported association between Rb and preservation of macrophage viability within otherwise oxidative stress environments such as TME (11). Indeed, gene expression analysis in AP-3-84-treated cells revealed the induction of several stress pathways once Rb was targeted including ER stress and an oxidative stress response. Gene enrichment analysis found two main cell death programs involved: p53-dependent and mitochondria-related pathways. These findings were supported by the loss of activity of AP-3-84 in the Rb/p53 myeloid conditional KO and by the detected changes in wild-type TAMs in the expression of apoptosis and p53-related proteins, ROS induction and cardiolipin accumulation. As described in previous studies, the combination of elevated ROS and ER stress/mitochondria crosstalk can induce cell death in macrophages (46,47). However, those pathways were redundant in our system as switching off only p53 signaling in p53 knock-out animals or using various pathway-specific inhibitors could not completely cancel AP-3-84-induced cell death as predicted by our gene expression data. However, we did demonstrate that targeting the LxCxE pocket of Rb is not redundant with the effects of clinically available CDK4/6 inhibitor palbociclib or HDAC inhibitors allowing further pre-clinical studies to investigate combination strategies. Specifically, it remains to be determined whether a combination between AP-3-84, palbociclib, HDAC inhibition, or ICI may provide a superior anti-tumor regimen.

Overall, we identify the Rb cleft pocket in M2 TAMs as a novel target for immunotherapy in ovarian cancer. The relationship between macrophage Rb expression in TME and cancer progression further supports that targeting the LxCxE cleft pocket of Rb in myeloid cells may provide a novel therapy strategy for TAM depletion in ovarian cancer patients.

## MATERIALS AND METHODS

### Experimental design

All studies were designed considering the estimates for sample size and power calculations to obtain statistically reliable data and with the help of the Wistar Institute Bioinformatics Facility when needed. All experiments were replicated two to five times and critical observations were made with different cell lines and models.

### Mice

All mice were maintained in a specific pathogen–free facility. C57BL/6 Albino mice (#000058), p53 knock-out mice (#002101), and E2F1 knock-out mice (#014531) were purchased from Jackson Laboratory. All experiments were approved by the Institutional Animal Care and Use Committee of the Wistar Institute. All efforts were made to minimize mouse suffering.

### Isolation of peritoneal thioglycolate-induced macrophages

Naïve C57BL6 mice were injected with 1 ml of 3% Thioglycollate sterile autoclaved solution in water. 4 days later peritoneal washes were collected and F4/80+ macrophages were sorted with magnetic beads (Miltenyi Biotec, Bergisch Gladbach, Germany) with the typical purity yield >95%.

### AP-3-84 sensitivity assay by flow cytometry, LDH and AlamarBlueHS assays

To check the sensitivity of different cell types to AP-3-84 cells were plated at 25 000/well in 96 U-bottom 96 well plate and cultured with different concentrations of AP-3-84 overnight for 16- 18 hours. Then the cells were washed and stained with the population-specific markers and Annexin V/DAPI staining with the following flow cytometry analysis. Peritoneal, splenic and ovarian cancer ascites macrophages (CD11b+ F4/80+), splenic CD4+ and CD8+ T cells (both pre-gated on CD3+ population) as well as different tumor cancer cell lines were analyzed. The percentage of Annexin V+ cells was measured and compared in the cultures with/without AP-3- 84 treatment.

Cell viability LDH and AlamarBlueHS assays were conducted according to the manufacturer’s protocol.

To assess the effect of inhibitors specific for various cell death pathways early in the cell death induction sequence, we applied a short-termed treatment with AP-3-84 in the presence of inhibitors with the following flow cytometry analysis. Specifically, we pre-incubated thioglycolate macrophages with the specified inhibitors of various cell death pathways (10 µM Nec-1s, 10 µM Z-Vad-FMK or 15 µM Ferrostatin-1) or three main ER stress pathways (20 µM BI09, 10 µM Ceapin-A7 and 5 µM GSK2656157 as respective IRE1α, ATF6 and PERK inhibitors) for 30 mins, then added 10 µM AP-3-84 and incubated cells for 40 more mins, washed the cells twice with cell culture media, added the specified inhibitors again and incubated the cells for 3 additional hours with the following Annexin V/DAPI staining, and flow cytometry analysis as described above. Increment of cell death was calculated as the difference of Annexin V+ frequencies between AP-3-84 treated and non-treated cells. Inhibition of cell death was calculated as the ratio of cell death increments in cells cultured with vs without inhibitor.

### Compound effects on tumor cell proliferation

To assess compound effects on tumor cell proliferation, tumor cells were stained with 5 uM CFSE and cultured in the presence of different concentrations of the tested compound (AP-3-84, palbociclib) for 13, 37 and 62 hours with the following flow cytometry analysis CFSE mean fluorescent intensity was measured and compared between different treatment conditions.

### In vivo ID8 ovarian cancer model with treatments

1.5×10^6^ ID8 ovarian cancer cells expressing luciferase (ID8-Luc cells) were injected intraperitoneally into B6 albino mice (B6(Cg)-Tyr c-2J /J, purchased from the Jackson Laboratory) or NSG mice. Around day 24 mice were injected with 3 mg luciferin in 200 ul PBS and tumor burden was assessed by bioluminescence measurements with in vivo imaging system (IVIS, PerkinElmer). Mice were split into groups based on the similar average bioluminescence signal between the groups. The specified treatments were run with the consecutive weekly measurements of tumor burden by IVIS until the end of the experiment. Mice were sacrificed and injected with 5 ml cold MACS buffer (PBS with 2 mM EDTA and 0.5% FCS) intraperitoneally. Then the peritoneal wall was opened by small cut, and cell suspension was collected for analysis.

For AP-3-84 treatments 7 daily injections with 5 mg/kg AP-3-84 were administered intraperitoneally starting from day 27.

For T cell depletion mice were treated with 200 ug/injection of anti-CD4 and anti-CD8 antibodies (BioXcell) twice per week starting simultaneously with AP-3-84 treatment on day 27. For palbociclib treatment mice were treated with 100 mg/kg palbociclib daily starting from day 27.

### Isolation of ID8-induced ascites macrophages

Peritoneal washes from ID8-bearing mice were collected on days 41-47 after ID8 tumor cell injection as specified and F4/80+ macrophages were sorted with magnetic beads (Miltenyi Biotec, Bergisch Gladbach, Germany) with the typical purity yield >95%.

### Measurements of tumor-specific responses in ID8-OVA ovarian cancer model

To assess the effects of AP-3-84 treatment on T cell tumor-specific responses we applied ID8- Luc cells overexpressing chicken ovalbumin protein (ID8-OVA cells). Similarly, to ID8 model 1.5×10^6^ ID8-OVA ovarian cancer cells were injected intraperitoneally into B6 albino mice, split into the groups with similar average tumor burden on day 24 and treated with 7 daily 5 mg/kg AP-3-84 injections. One week after the end of the treatment mice were sacrificed. Splenocytes and ascitic cells were isolated and re-stimulated with SIINFEKL peptide (immunodominant ovalbumin peptide) for 12 hours in the presence of 2 uM Monensin (BioLegend, San Diego, CA) with the following intracellular staining of IFN-γ and measurement of IFN-γ-producing CD8+ T cells (meaning ovalbumin-specific T cells) by flow cytometry.

### Generation of M1 and M2 type polarized macrophages

Bone marrow was collected from femur and tibia of C57BL6 mice. Hematopoietic progenitor cells (HPC) were collected using Lineage Cell Depletion kit (Miltenyi Biotec, Bergisch Gladbach, Germany or R&D Systems, Minneapolis, MN). Then HPC were cultured in complete RPMI 1640 media with 10 ng/mL M-CSF (PeproTech, Cranbury, NJ) for 6 days at 37°C and 5% CO2. Culture media were changed every 3 days. Cells were incubated for additional 24 hours with LPS (100 ng/mL) and murine IFN-γ (50 ng/mL) for M1 type polarization or murine IL-4 (10 ng/mL) and murine IL-13 (10 ng/mL) for M2 type polarization, respectively. M1 or M2 polarized murine macrophages were treated with AP-3-84 for 16 hours, collected and stained with Annexin V (BioLegend, San Diego, CA) and Live/Dead Aqua (Thermo Fisher Scientific, Waltham, MA). To measure *Rb1* gene expression, either M1 or M2 murine macrophages were collected with the following conducted RT-PCR analysis for Rb1 and 18S genes. For cytoplasmic staining of Rb protein, cells were fixed and permeabilized with Cytofix/Cytoperm kit (BD, Franklin Lakes, NJ) and then stained with anti-Rb antibodies and their corresponding secondary IgG (#MA5-11387 Thermo Fisher Scientific, Waltham, MA, #NB100-56598 Novus Biologicals, Centennial, CO).

### Flow cytometry

Ascites cells were collected by peritoneal washes as described above, centrifuged at 400 x g, 5 min, +4°C, resuspended in 1 ml cold ACK lysis buffer to lyse red blood cells, washed once with 10 ml cold MACS buffer (0.5% FBS, 2 mM EDTA in PBS) with the following 400 x g, 5 min, +4°C centrifugation and about 1 x 10^6^ cells were dispensed for flow staining in 96 U-bottom well plate. Plates were centrifuged as above and supernatants were removed. Cells were resuspended by mild vortexing and 50 ul of 2.5 µg/ml final (meaning 1/200 dilution of the stock) Fc-block (anti-mouse anti-CD16/32 antibodies, TruStain FcX from Biolegend, San Diego, CA) were added with the following addition of 1/200 dilution of antibodies for surface marker staining and Aqua Live/Dead dye (Thermo Fisher Scientific, Waltham, MA). Cells were incubated 40 minutes at +4°C, washed once with MACS buffer, fixed in 1% paraformaldehyde in PBS and analyzed by BD FACSymphony instrument and FlowJo software (BD Biosciences). For intracellular staining cells were first stained for surface markers as described above, then washed with MACS buffer, permeabilized with BD Cytofix/Cytoperm kit (BD Biosciences) for 20 minutes at +4°C, incubated with the specified antibodies for intracellular markers for 40 minutes, washed once with MACS buffer and fixed in 1% paraformaldehyde in PBS.

For reactive oxygen species (ROS) measurement attached macrophages were loaded with 10 µM CM-H_2_DCFDA in PBS for 30 minutes at 37°C and 5% CO_2_, washed twice with cell culture media and treated with 10 µM AP-3-84 for 20 minutes. Then the cells were washed twice with cell culture media and incubated in cell culture media for 2 hours more with the following flow cytometry analysis of ROS-induced fluorescence in FITC-channel (excitation with 488 laser and detection in 515/20 filter channel).

### Western Blotting

For Western blotting whole cell lysates were fractionated by 10% Bis-Tris polyacrylamide NuPAGE gels (Thermo Fisher Scientific, Waltham, MA) and transferred to a polyvinylidene difluoride membrane using a transfer apparatus according to the manufacturer’s protocols (Bio-Rad, Hercules, California). After blocking with 5% nonfat milk in PBST (PBS, pH 7.5, 0.5% Tween 20) for 60 min, the membrane was washed once with PBST and incubated overnight at +4°C with the specified primary antibody, washed 3 times, 5 minutes/wash, with PBST and incubated for 1 h at room temperature the corresponding secondary antibody with the following ECL development (Amersham Biosciences).

For protein analysis in ascites macrophages, ascites macrophages from ID8 bearing mice (day 45) were isolated and treated with the specified AP-3-84 concentrations or for the specified time. Then cells were washed and lysed, and the expression of the apoptosis-related proteins was measured by western blot.

### Immunofluorescence (IF) staining

For ascites confocal microscopy analysis, single cells from ID8 tumor bearing mice were seeded on serum-coated cover glass (20 mm round cover glass from CELLTREAT Scientific Products, Pepperell, MA), cultured 48 hours with complete RPMI 1640 with 10% FBS (Thermo Fisher Scientific, Waltham, MA), 50 U/mL Penicillin and 50 ug/mL streptomycin (Millipore Sigma, Burlington, MA). For BMDM confocal microscopy, hematopoietic progenitor cells (HPC) were isolated as described above and seeded onto cover glass, then cultured for 5 days with M-CSF and an additional 1 day with LPS/IFN-γ or IL-4/IL-13 for M1 or M2 polarization, respectively, as mentioned above for BMDM generation. After washing with PBS, cells were fixed in 4% paraformaldehyde (Electron Microscopy Sciences, Hatfield, PA), permeabilized, then blocked in 1% BSA/PBS for 1 hour at room temperature. Primary antibodies, anti-Rb (clone 1F8, 1:100 dilution, Invitrogen, Waltham, MA), anti-F4/80 (clone ab6640, 1:100 dilution, Abcam, Cambridge, UK) and anti-E2F1 (clone KH95, 1:100 dilution, Invitrogen) were incubated for 16 hours at 4°C. After washing with PBS, cells were incubated with corresponding secondary antibodies (anti-Mouse conjugated to AF488 and anti-Rat conjugated to Cy5), DAPI and Phalloidin conjugated with AF594 (for F-actin staining, Thermo Fisher Scientific, Waltham, MA) for 1 hour at room temperature, then mounted on slide glass in VECTASHIELD antifade mounting media (Vector Laboratories, Newark, CA). Confocal images were obtained using Leica TCS SP8 confocal microscope (Leica Mycrosystems, Wetzlar, Germany).

### RNA-Sequencing

3’ mRNA-Seq libraries were generated from 100 ng of DNAse I treated total RNA using the QuantSeq FWD Library Preparation kit (Lexogen, Vienna, Austria), according to the manufacturer’s directions. Overall library size was determined using the Agilent Tapestation and the DNA 5000 Screentape (Agilent, Santa Clara, CA). Libraries were quantitated using real-time PCR (Kapa Biosystems, Wilmington, MA). Libraries were pooled and Single read, 75 base pair Next Generation Sequencing was done on a P2 flow cell using the NextSeq 2000 (Illumina, San Diego, CA).

### RNA-Sequencing analysis

RNA-seq data was aligned using the STAR (48) algorithm against the mm10 mouse genome version and RSEM v1.2.12 software (49) was used to estimate read counts and RPKM values using gene information from mouse Ensemble transcriptome version GRCm38.89. Raw counts were used to estimate significance of differential expression difference between two experimental groups using DESeq2 (50). Overall gene expression changes were considered significant if they passed FDR<5% threshold unless stated otherwise. Gene set enrichment analysis was done using QIAGEN’s Ingenuity® Pathway Analysis software (IPA®, QIAGEN Redwood City, www.qiagen.com/ingenuity) using “Canonical pathways” and “Upstream Regulators” options. Top results that passed FDR<5% threshold and had a significantly predicted activation state (|Z|>1) were reported. Targeted cell death related enrichment analysis was performed with single sample GSEA (ssGSEA) algorithm (51) using MSigDB (32) categories containing keywords CELL_DEATH, NECROSIS, NECROTIC, APOPTOSIS, APOPTOTIC, AUTOPHAGY, FERROPTOSIS. Samples were scored by ssGSEA and compared between groups using R package limma (52). Results that pass p<0.05 thresholds were considered and reported p53 and mitochondria-related enrichment results were derived from MSigDB gene sets WU_APOPTOSIS_BY_CDKN1A_VIA_TP53 and GOBP_NEGATIVE_REGULATION_OF_MITOCHONDRIAL_OUTER_MEMBRANE_PER MEABILIZATION_INVOLVED_IN_APOPTOTIC_SIGNALING_PATHWAY correspondingly. Macrophage Principal Component Analysis was based on 305 genes most significantly differentially expressed between M0-M1-M2 macrophages in p53 wild-type condition. Average gene expression across replicates was calculated, producing a 305 gene signature for each M0/M1/M2 macrophage type. Correlation coefficients using the three 305 gene signatures were estimated for each sample and z-scores were calculated across the 3 correlation coefficients used in principal component analysis.

### TCGA analysis

TOIL RSEM transcript per million (TPM) Gene expression data was downloaded from the Xena browser (https://xenabrowser.net) (53). Gene expression values were correlated across samples using Spearman rank correlation.

Gene expression values between healthy tissues and tumors were compared using a one-sided Wilcoxon rank-sum test.

Kaplan Meier survival curves were generated by splitting patients into high and low expression groups, using the median gene expression levels as threshold, where the log-rank p-value is used for survival analyses.

To quantify immune cells, CIBERSORT software (54) was applied to TCGA samples using the default set of 22 immune-cell signatures.

### Human Protein Atlas Data

Human Protein Atlas Data for the Rb1 gene expression dataset based on summarized meta-analysis of single-cell RNA sequencing of various human samples was downloaded from https://www.proteinatlas.org/about/download website, dataset #8. Transcript expression levels summarized per gene in different immune cells from various tissues are graphed as normalized expression (“nTPM”) as obtained from the website.

### Lipidomics analysis by LC-MS/MS

Thioglycolate-elicited or ID8 ascites macrophages were isolated by magnetic F4/80 sorting, seeded 3.5 x 10^5^ cells/well into 6 well plates and rested overnight in cell culture media. Then the cells were treated with 10 uM AP-3-84 for 4 hours, washed twice with PBS and collected with the cell scraper for lipidomics analysis.

LC/ESI-MS analysis of lipids was performed on a Thermo HPLC system coupled to an Thermo Scientific™ Orbitrap Fusion™ Lumos™ Tribrid™ mass spectrometer. Phospholipids were separated on a normal phase column (Luna 3 μm Silica (2) 100 A, 150 × 1.0 mm, (Phenomenex)) at a flow rate of 0.065 ml/min. The column was maintained at 35 °C. The analysis was performed using gradient solvents (A and B) containing 10 mM ammonium formate. Solvent A contained isopropanol/hexane/water (285:215:5, v/v/v), and solvent B contained isopropanol/hexane/water (285:215:40, v/v/v). All solvents were LC/MS-grade. The gradient was as follows: 0-3 min, 10-37% B; 3-15 min, hold at 37% B; 15-23 min, 37-100% B; 23-75 min, hold at 100% B; 75-76 min, 100-10% B; 76-90 min, equilibrate at 10% B. Lipids were analyzed in negative ion mode using the following parameters: capillary voltage, 3500; sheath, aux and sweep gases (35, 17, 0, respectively); ion transfer tube temperature, 300°C; orbitrap resolution, 120,000; scan range 400-1800 m/z; Rf lens, 40; injection time, 100 ms; intensity threshold set at 4e^2^. For data dependent MS^2^, an isolation window of 1.2 m/z was used. Collision energy (HCD) was static at 24 with an orbitrap resolution of 15,000 with an injection time of 22 ms. Lipids and their oxygenated metabolites were identified with high accuracy by exact masses using Thermo Scientific™ Orbitrap Fusion™ Lumos™ Tribrid™ mass spectrometer and confirmed by MS^2^ analysis. Compound Discoverer^TM^ software package (Thermo Fisher Scientific, Waltham, MA) with an in-house generated analysis workflow and oxidized phospholipid database was used. Lipid signals (with a signal/noise ratio of >3) were further filtered by retention time and values for *m/z* were matched within 5 ppm to identify the lipid species. 1-stearoyl-2-oleoyl-sn-glycero-3-phosphoethanolamine and 1-stearoyl-2-arachidonoyl-sn-glycero-3-phosphoethanol-amine, 1-stearoyl-2-oleoyl-sn-glycero-3-phospho-L-serine (sodium salt), 1-stearoyl-2-oleoyl-sn-glycero-3-phosphocholine, 1-palmitoyl-2-oleoyl-sn-glycero-3- phospho-inositol (ammonium salt), 1,1’,2,2’-tetralinoleyl-cardiolipin (sodium salt) (Avanti Polar Lipids) were used as reference standards to build the calibration curves. Deuterated phospholipids: 1-hexadecanoyl(d31)-2-(9Z-octadecenoyl)-sn-glycero-3-phospho-ethanolamine, 1-hexadecanoyl(d31)-2-(9Z-octadecenoyl)-sn-glycero-3-phosphocholine, 1-hexadecanoyl(d31)- 2-(9Z-octadecenoyl)-sn-glycero-3-phosphoserine, 1-hexa-decanoyl(d31)-2-(9Z-octadecenoyl)-sn-glycero-3-phosphate, 1-hexadecanoyl(d31)-2-(9Z-octadecenoyl)-sn-glycero-3- phosphoglycerol, 1-hexadecanoyl(d31)-2-(9Z-octadecenoyl)-sn-glycero-3-phospho-(1’-myo-inositol) and 1,1’,2,2’-tetramyristoyl-cardiolipin (sodium salt) (Avanti Polar Lipids) were used as internal standards. Internal standards were added directly to the MS sample to a final concentration of 1 µM.

### Patient samples

Patient samples were obtained under the Christiana Care Institutional Review Board approval of protocol CCC#41068 entitled *Evaluation of Immune Response by a Novel Small Molecule RB Protein Modulator in Patients with blood and solid cancer.* Informed consent was obtained after the nature and possible consequences of the studies were explained. Ascitic fluid was collected prior to surgery or at the time of the debulking surgery or prior to neoadjuvant treatment. Ascites cells were treated with AP-3-84 or vehicle for 16 hours with the following Annexin V+ cell frequencies analysis in CD45+CD11b+CD163+CD68+ macrophages and CD3+ T cells by flow cytometry.

### General Methods for Chemistry

All commercially obtained solvents and reagents were used as received. Flash column chromatography was performed using silica gel 60 (230-400 mesh). Analytical thin layer chromatography (TLC) was carried out on Merck silica gel plates with QF-254 indicator and visualized by UV, PMA, or KMnO4. ^1^H NMR spectra were recorded on a Bruker Advance 400. Chemical shifts are reported in parts per million (ppm, δ) using the residual solvent line as a reference. Splitting patterns are designated using the following abbreviations: s, singlet; d, doublet; t, triplet; dd, doublet of doublet; m, multiplet; br, broad. Coupling constants (J) are reported in hertz (Hz). Tetramethylsilane was used as an internal standard for proton nuclear magnetic resonance for samples run in CDCl3 or DMSO-d6. LC−MS data were acquired on a Waters Acquity UPLC/MS system equipped with a UPLC binary pump, an SQD 3100 mass spectrometer with an electrospray ionization (ESI) source and a PDA detector (210−400 nm). High-resolution mass spectra were obtained using the Q Exactive HF-X mass spectrometer which provided high resolution, accurate mass, and total ion and extracted ion chromatograms. All compounds tested were present within a 5 ppm mass error. The purity of all final compounds was determined by HPLC, and the compounds are at least ≥95% pure.

### Synthesis of 2-(3-Methoxyphenyl)-4-methyl-1,2,4-thiadiazolidine-3,5-dione (AP-3-84)

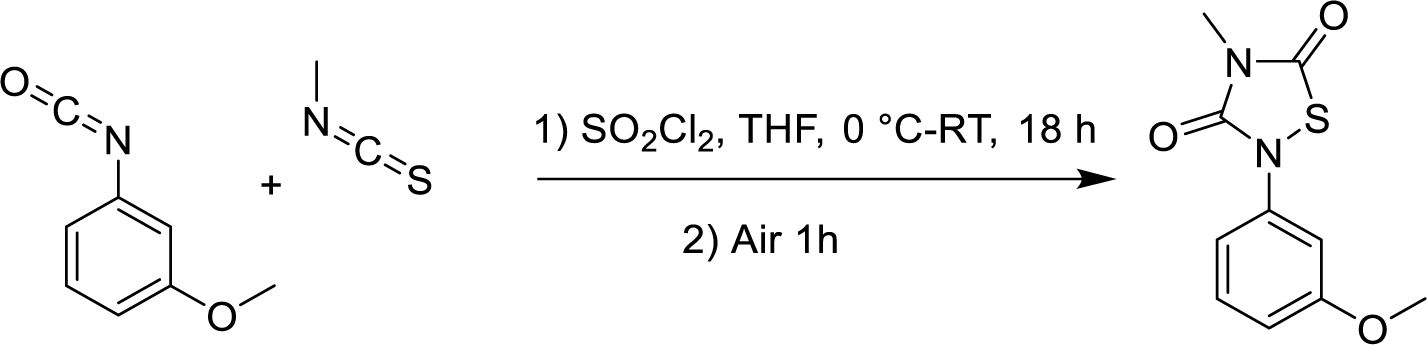

A stirred solution of 1-isocyanato-3-methoxybenzene (4.0 g, 27.56 mmol) and Isothiocyanatomethane (2.01 g, 27.56 mmol) in THF (60 mL) was cooled to 0 °C. Sulfuryl chloride (3.72 g, 27.56 mmol) was added, and the mixture was allowed to warm to room temperature and stirred overnight. The reaction was then opened to the air and stirred for 30 minutes before the solvent was removed under reduced pressure. Crude product passed through the silica gel with 20 % ethyl acetate in hexane. The product precipitated in methanol and washed with cold methanol filter solid dried under reduced pressure afforded the title compound (2.0 g, 8.39 mmol). Product was confirmed by LC-MS and ^1^H NMR.

^1^H NMR (400 MHz, CDCl3) δ 7.33 (dd, *J* = 13.7, 5.5 Hz, 1H), 7.14 (t, *J* = 2.3 Hz, 1H), 7.05 (ddd, *J* = 8.0, 2.1, 0.8 Hz, 1H), 6.83 (ddd, *J* = 8.4, 2.4, 0.7 Hz, 1H), 3.83 (s, 3H), 3.29 (s, 3H).

LC-MS calculated C_10_H_11_N_2_O_3_S^+^ [M + H] ^+^ 238.04, Found 239.01.

### Synthesis of 2-(3-(3-Aminopropoxy)phenyl)-4-methyl-1,2,4-thiadiazolidine-3,5-dione (JS- 102)

JS-102 was synthesized at Medinoah Inc., a company specializing in custom synthesis. The spectral characterization data is provided.

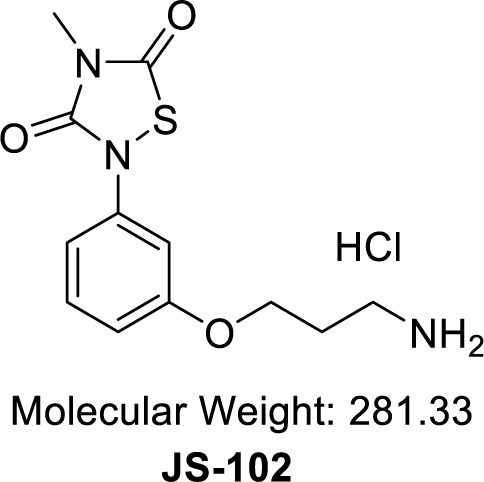

^1^H NMR (400 MHz, DMSO) δ 8.25 (s, 3H), 7.38 (t, *J* = 8.2 Hz, 1H), 7.24 – 7.14 (m, 1H), 7.10 (dd, *J* = 7.8, 1.8 Hz, 1H), 6.92 (dd, *J* = 8.3, 2.3 Hz, 1H), 4.10 (t, *J* = 6.1 Hz, 2H), 3.13 (s, 3H),

2.95 (dd, *J* = 12.6, 6.6 Hz, 2H), 2.18 – 1.97 (m, 2H).

LC-MS Calculated C_12_H_15_N_3_O_3_S [M + H] ^+^ 281.08, Found 281.88.

### Synthesis of 1-(3’,6’-Dihydroxy-3-oxo-3H-spiro[isobenzofuran-1,9’-xanthen]-5-yl)-3-(3-(3- (4-methyl-3,5-dioxo-1,2,4-thiadiazolidin-2-yl)phenoxy)propyl)thiourea (AP-08-239)

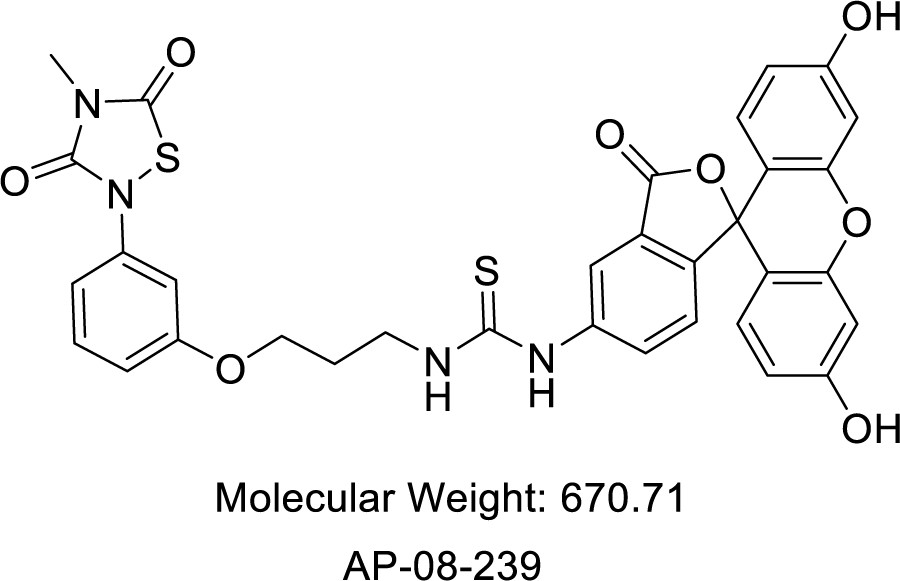

To a stirred solution of 2-(3-(3-aminopropoxy)phenyl)-4-methyl-1,2,4-thiadiazolidine-3,5-dione (50 mg, 0.16 mmol) in 3 mL of dry DMF at 0 °C was simultaneously added FITC (61 mg, 1.6 mmol) and pyridine (24.9 mg, 0.32 mmol). The reaction mixture was slowly brought to room temperature and stirred for 16h. Completion of the reaction was confirmed by LC-MS, and the crude reaction mixture was purified by reverse phase HPLC to afford the title compound as an orange-yellowish solid. (40 mg, 0.059 mmol). Product was confirmed by LC-MS and ^1^H NMR. ^1^H NMR (400 MHz, MeOD-D4) δ 8.17 (s, 1H), 7.81 (d, *J* = 7.1 Hz, 1H), 7.33 (t, *J* = 8.2 Hz, 1H), 7.24 (t, *J* = 2.2 Hz, 1H), 7.18 (d, *J* = 8.3 Hz, 1H), 7.06 (dd, *J* = 8.0, 1.5 Hz, 1H), 6.98 – 6.71 (m, 5H), 6.69 (d, *J* = 7.7 Hz, 2H), 4.14 (t, *J* = 6.0 Hz, 2H), 3.9-2.77 (m, 2H), 3.20 (s, 3H), 2.25-2.05 (m, 2H).

LC-MS calculated C_33_H_26_N_4_O_8_S_2_ [M + H] ^+^ 670.71, Found 671.21.

### Protein Expression and purification

GST-Rb 379-928 was expressed and purified by Biologics Corp (Indianapolis, IN). To generate untagged Rb, this expression construct was then cloned into a HIS-SUMO expression vector by Epoch Biosciences. The construct was transformed in T7 Express E coli (NEB) and grown to O.D. of 0.8 in TB media at 37 C. The culture was then induced with 1 mM IPTG and grown overnight at 17C with constant shaking. The culture was then centrifuged at 5000 x g for 10 min. The pellets were resuspended in HIS purification buffer (50 mM KPO4, pH 8.0, 250 mM NaCl, 1 mM MgCl2, 10 mM imidazole, 10% glycerol, 0.05% Triton X100) with protease inhibitor cocktail. After resuspension, 1 mg/mL lysozyme and 1 ug/mL DNaseI were added and the mixture sonicated 3 times for 30s on ice at setting 6 in between 10-15 min intervals. After sonication, the mixture was centrifuged at 26,000 x g for 20 min at 4C to pellet insoluble material. The supernatant was passed through a 1mL column packed with TALON cobalt resin which had been equilibrated with HIS purification buffer. The column was then washed with 40 mL of HIS-purification buffer and the protein eluted in 5 x 1 mL fractions of HIS-purification buffer supplemented with 100 mM imidazole. The fractions were pooled and dialyzed overnight against HIS-purification buffer in the presence of 25 ug/mL HIS-SUMO protease to remove the HIS-SUMO tag and reduce the imidazole concentration. After dialysis, the protein was passed through the 1 mL TALON cobalt resin column again to bind the free HIS-SUMO tag and the HIS-SUMO protease and the flow-through contained untagged-Rb. Protein was aliquoted and stored at -80 C.

HIS-SUMO-MDM2. HIS-E7 1-97, and HIS-SUMO-C-A mutant Rb 380-785 were expressed and purified using the same methods as above. HIS-E7 1-97 was synthesized and cloned into pET 28a+ expression vector at Genscript. The other two constructs were prepared at Epoch Biosciences. The MDM2 construct was cloned into the HIS-SUMO expression vector from pGEX-4T MDM2 WT, which was a gift from Mien-Chie Hung (Addgene plasmid # 16237 ; http://n2t.net/addgene:16237 ; RRID:Addgene_16237). HIS-E7 1-97 was induced at 20 C instead of 17 C like the HIS-SUMO constructs. The resulting elutions were concentrated, aliquoted and stored at -80 C. We did not dialyze with SUMO protease since we needed the HIS tag for downstream assays in this case.

The constructs containing the GST-tagged Rb fragments were made by Epoch Biosciences. Expression and purification of these proteins were similar to the above, except the induction temperature was 20 C. The GST purification buffer was 100 mM Tris, pH 8.0, 250 mM NaCl, 10% glycerol, 0.05% TX100. The soluble fraction was passed through a 1 mL glutathione resin (Thermo Scientific) column and the column was washed with 40 mL of GST purification buffer. Fractions were eluted from the column with GST purification buffer containing 10 mM glutathione. Fractions were pooled, concentrated in aliquoted and stored at -80 C.

E2F-1 wt-pGex2TK was a gift from William Kaelin (Addgene plasmid # 21668 ; http://n2t.net/addgene:21668 ; RRID:Addgene_21668). The construct was cloned into N-terminal maltose-binding protein (MBP) expression vector at Epoch Biosciences. Expression and purification were similar to what was described for other proteins, except the induction temperature was 20 C and the soluble fraction was passed through a 1 mL column packed with amylose resin (Thermo Scientific). The purification buffer was the same as what we used for the GST-protein purification. The fractions were eluted in purification buffer containing 10 mM maltose, then pooled, concentrated, aliquoted and stored at -80C.

HDAC1-FLAG and HDAC4-FLAG were purchased from Active Motif (Cat#: 31504 and Cat#: 31527, respectively).

### AP-08-239 binding to Rb by Homogeneous Time Resolved Fluorescence (HTRF)

All assays were conducted in 10 uL in white, opaque, low volume 384-well plates in assay buffer (10 mM HEPES, pH 7.4, 100 mM NaCl, 5 mM MgCl2, 0.01% Triton X100). GST-Rb (379–928) or HIS-SUMO-C-A mutant Rb 380-785 was pre-incubated with 0.4 nM HTRF donor conjugated antibodies anti-GST-Tb or anti-HIS-Tb (PerkinElmer), respectively, at 2X the final concentration as indicated on the figures. Assays were initiated by the addition of 5 uL 2X AP- 08-239 as indicated on the figures. For compound inhibition experiments, the GST-Rb 379-928 concentration was 5 nM. Test compounds were added to 5 uL assays containing GST-Rb/anti-GST-Tb by direct dilution using the Echo 650 acoustic liquid handling system in 100 nL of DMSO before the addition of 30 nM AP-08-239 (15 nM final). After 30 min, the HTRF signal was measured using a ClarioStar plate reader (BMG LabTech) with an excitation wavelength of 320 nm, emission wavelengths of 520 and 620 nm with a 50 us delay and 200 us assay window. The HTRF ratio was then determined by dividing the RFU at 520 nm by the RFU at 620 nm with a 10,000 multiplier. For compound inhibition experiments, % inhibition values were calculated from the HTRF ratios such that 0% is the HTRF ratio in the absence of test compound, and 100% is the HTRF ratio in the absence of Rb protein.

### AP-08-239 binding to untagged Rb by Fluorescence Polarization (FP)

Rb 379-928 was serially diluted from 0 to 500 nM in assay buffer and 5 uL was dispensed into a black, opaque, low volume, 384-well plate. Assays were initiated by the addition of 5 uL of 40 nM AP-08-239 (20 nM final concentration). After 30 min, the FP signal was measured using an Envision plate reader (PerkinElmer). mP was calculated using the following formula, mP =[(S - P*G)/(S + P*G)]*1000, where S is the parallel fluorescence intensity, P is the perpendicular fluorescence intensity, and the G factor set to 0.95.

### AP-03-084 inhibition of GST-Rb protein-protein interactions

HTRF assays were developed for GST-Rb 379-928 binding to either HIS-E7, HIS-SUMO- MDM2, MBP-E2F1, and HDAC1-FLAG using the same assay buffer as described above. Each protein was preincubated with 0.4 nM of its corresponding HTRF donor or acceptor antibody prior to being added to the assay. All antibodies were obtained from PerkinElmer Life Sciences except for the anti-FLAG-FITC conjugated antibody which was purchased from BioLegend, San Diego, CA. For HIS-E7 (10 nM) and HIS-SUMO-MDM2 (5 nM), anti-HIS-d2 HTRF acceptor antibody was used, and anti-GST-Tb HTRF donor antibody was used for GST-Rb379-928 (5 nM). For MBP-E2F1 (5 nM), anti-MBP-Tb HTRF donor antibody was used, and anti-GST-d2 acceptor antibody was used with GST-Rb 379-928 (5 nM). For HDAC1-FLAG and HDAC4-FLAG (5 nM), anti-FLAG-FITC conjugated HTRF acceptor antibody was used, and anti-GST-Tb HTRF donor antibody was used for GST-Rb 379-928 (5 nM). Titration curves of test compounds were added in 100 nL DMSO by direct dilution using the Echo 650 acoustic liquid handling system to 5 uL of GST-Rb 379-928. After a 15 min pre-incubation, the 5 uL of the other binding protein was added. After 1 hour, the HTRF signal was measured using the ClarioStar plate reader. For d2 HTRF acceptors, the emission wavelengths were 665 nm and 620 nm, and for the FITC HTRF acceptor, the emission wavelengths were 520 nm and 620 nm. The HTRF ratios and % inhibition values were calculated as described above.

### AP-03-084 displacement of GST-Rb protein-protein interactions

For displacement assays, the assays were set up as described above, except the titration curve of AP-03-084 was added to a pre-formed complex of GST-Rb379-928 and the corresponding interacting protein (MBP-E2F1, HIS-SUMO-MDM2, or HDAC1-FLAG). After 1 hr, the HTRF signal was measured. The HTRF ratios and % displacement values were calculated as described above, where 0% displacement is the HTRF ratio in the absence of test compound, and 100% displacement is the HTRF ratio in the absence of GST-Rb.

### HDAC binding to GST-Rb protein fragments

To determine where HDAC1 bound to Rb, GST-tagged Rb fragments 379-928, 380-785, 380- 600, and 785-928 were expressed and purified as described above. 10 nM of GST-Rb fragment was prebound to anti-GST-Tb HTRF donor antibody and then incubated with either 10 nM or 1 nM HDAC1 FLAG prebound to anti-FLAG FITC conjugated HTRF acceptor antibodies. After 1 hour, the HTRF signal was measured. To test if AP-03-084 could inhibit HDAC1-FLAG binding to the GST-Rb fragments, titration curves of AP-03-084 in 100 nL 100% DMSO was added 5 nM of each GST-Rb fragment prebound to HTRF antibody using the Echo 650 acoustic liquid handling system. After a 15 min pre-incubation, 5 nM HDAC1-FLAG was added to start the reaction and 1 hr later the HTRF signal and % inhibition values were calculated as described above.

### Statistical analysis

Statistical analysis was performed using unpaired two-tailed Student’s t-test with significance determined at 0.05. Estimation of variation within each data group was performed and variance was similar between groups that were compared. GraphPad and R/Rstudio software with ggplot2 package were used to graph the data and calculate the corresponding statistics.

The percentage of different cell populations with tumor burden was correlated by the Pearson method.

tSNE analysis of flow cytometry data was performed using built-in tSNE analysis function in FlowJo software (BD Biosciences).

## SUPPLEMENTARY MATERIALS

**Supplementary Fig. S1. Targeting the LxCxE cleft pocket of Rb protein with AP-3-84 modulates its interactions with the adaptor proteins.**

**Supplementary Fig. S2. Therapeutic myeloid Rb targeting preferentially affects macrophages, but not tumor cells and demonstrates the effects different from CDK4/6 palbociclib inhibitor.**

**Supplementary Fig. S3. Therapeutic myeloid Rb targeting changes the composition of immune cell populations in the ascites of ID8-bearing mice and preferentially affects M2 type polarized macrophages.**

**Supplementary Fig. S4. Myeloid Rb targeting initiates dramatic changes in macrophage gene expression profile and lipid composition inducing the major stress response pathways.**

**Supplementary Fig. S5. Therapeutic Rb targeting upregulates p53 expression in macrophages.**

**Supplementary Fig. S6. Therapeutic Rb targeting delays tumor growth in various ovarian cancer models and demonstrates the effects non-redundant to CDK4/6 palbociclib inhibitor.**

**Supplementary Fig. S7. Comparison of patient survival outcome with different expression levels of RBL1/RBL2 in tumor.**

**Supplementary Table S1. The effect of HDAC inhibitors on Rb-HDAC1 interaction.**

**Supplementary Table S2. M0/M1/M2 signature genes.**

## Data availability

All data in the figures are available in the published article and in online supplemental material. Unprocessed images of immunoblots in Figure 5 are submitted as Source File linked to that figure. RNA Sequencing raw data is submitted to GEO repository and can be accessed there (accession number GSE239479).

## Acknowledgements

We thank Drs. Dmitry Gabrilovich, Jose Conejo-Garcia, and Alfredo Perales-Puchalt for critical discussion.

## Authors contribution

E.N.T. participated in research design, performed most of the experiments, analyzed the results, wrote manuscript; T.K., X.Y. participated in research design, performed most of the experiments, analyzed the results; A.N.R.P. synthesized the chemical compounds, C.H., A.B, J.C, L.L, L.D, V.A.T., P.S., M.DG.C.M., D.Z., Y.Y.T., H.B. performed some experiments, analyzed the results; A.K., N.A. analyzed gene expression data and publicly available datasets on ovarian cancer patients; M.G.C., S.J., S.C.-P., B.B. provided clinical samples and analyzed the results; D.W., R.Z. participated in research design and provided financial support; V.E.K., J.M.S. participated in research design, wrote parts of the manuscript; L.J.M. obtained financial support for the study, designed overall concept and specific experiments, supervised experiments, interpreted results, and wrote manuscript.

## Supplementary Figures

**Supplementary Fig. S1.**
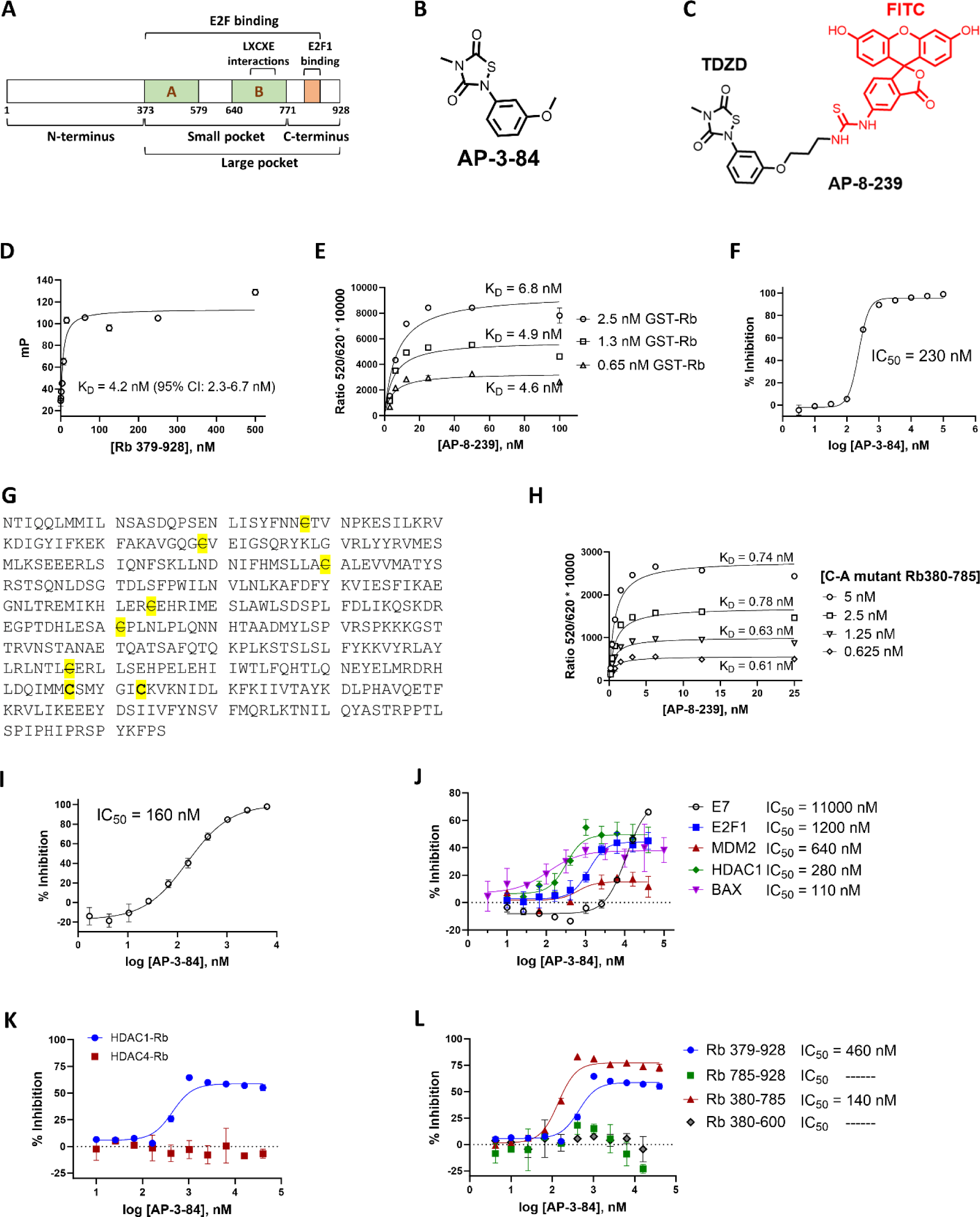
Targeting the LxCxE cleft pocket of Rb protein with AP-3-84 modulates its interactions with the adaptor proteins. **(A)** Rb protein scheme according to (18). **(B)** Chemical structure of AP-3-84. **(C)** Chemical structure of AP-8-239 (AP-3-84-FITC) probe. **(D)** AP-8-239 binds to untagged Rb (379–928), measured by fluorescence polarization (FP). **(E)** AP-8-239 binds to GST-Rb (379–928) as measured by HTRF. **(F)** Inhibition AP-8-239 binding to GST-Rb (379–928) by AP-3-84 compound. AP-3-84 was first pre-incubated with Rb, followed by AP-08-239 addition and HTRF measurement. **(G)** Rb (380–785) protein sequence. All the highlighted cysteines were mutated to alanine (indicated with a strikethrough font) except Cys706 and Cys712. **(H)** AP-8-239 binds to Cys706 and Cys712 of mutant Rb380-785, measured by HTRF as above. **(I)** AP-3-84 inhibits AP-8-239 binding to C-A mutant 380-785 measured by HTRF as above. **(J)** AP-3-84 inhibits the binding of Rb interacting proteins, measured by HTRF as above. **(K)** AP-03-84 inhibits HDAC1, but not HDAC4 binding to mutant GST-Rb379-928. **(L)** AP-3-84 inhibits HDAC1 binding to Rb by binding to the 380-785 region, measured by HTRF as above. Mean +/–SD are shown (n=2-4 per condition). Representative experiment out of at least 3 repeats is shown.

**Supplementary Fig. S2.**
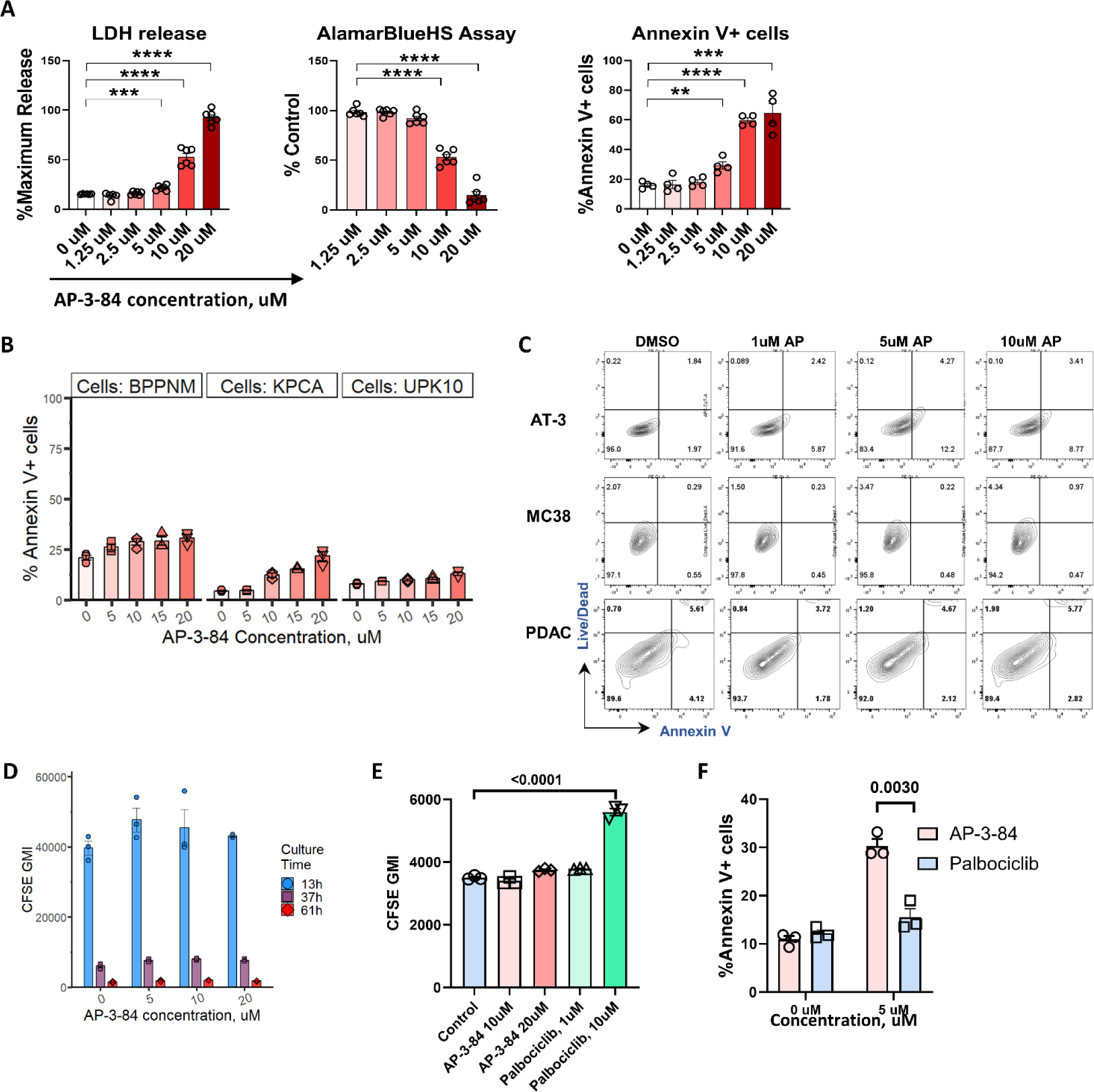
Therapeutic myeloid Rb targeting preferentially affects macrophages, but not tumor cells and demonstrates the effects different from CDK4/6 palbociclib inhibitor. **(A)** Thioglycollate-induced macrophages were isolated with magnetic F4/80-beads from the peritoneal washes of mice 4 days after thioglycolate injection. Macrophages were incubated 16-18 hours with different concentrations of AP-3-84. Viability of peritoneal macrophages was measured by LDH assay, AlamarBlueHS assay and Annexin V+/Live-Dead flow cytometry analysis. n=4-6 per condition. **(B)** AP-3-84 effect on viability of different ovarian cancer cell lines: BPPNM, KPCA, UPK10. Cell treatment and annexin V staining were conducted as in **Fig. 1B,C**. n=3 per condition. **(C)** AP-3-84 effect on viability of AT-3 mammary carcinoma, MC-38 colon cancer, and PDAC pancreatic cancer cells was tested as in (**C**) by Annexin V+/Live-Dead flow cytometry analysis. **(D)** ID8 tumor cell proliferation in the presence of AP-3-84 at specified concentrations was measured by CFSE dilution assay after 13h, 37h and 61h of ID8 cell coincubation with AP-3-84. The similar drop of CFSE geometric mean intensity (CFSE GMI) during the cell culture is shown. n=3 per condition. **(E)** The effect of 2-day treatment of ID8 tumor cells with the specified concentrations of AP-3- 84 and palbociclib on cell proliferation measured by CFSE dilution assay. n=3 per condition. **(F)** Ascites macrophages were isolated from ID8-bearing mice, incubated with the specified concentrations of AP-3-84 or palbociclib overnight and cell viability was measured by Annexin V/DAPI staining. n=3 per condition. Means +/– SEM are shown with indicated exact p-values or **p<0.01, ***p< 0.001, **** p<0.0001 by unpaired Student t-test comparison.

**Supplementary Fig. S3.**
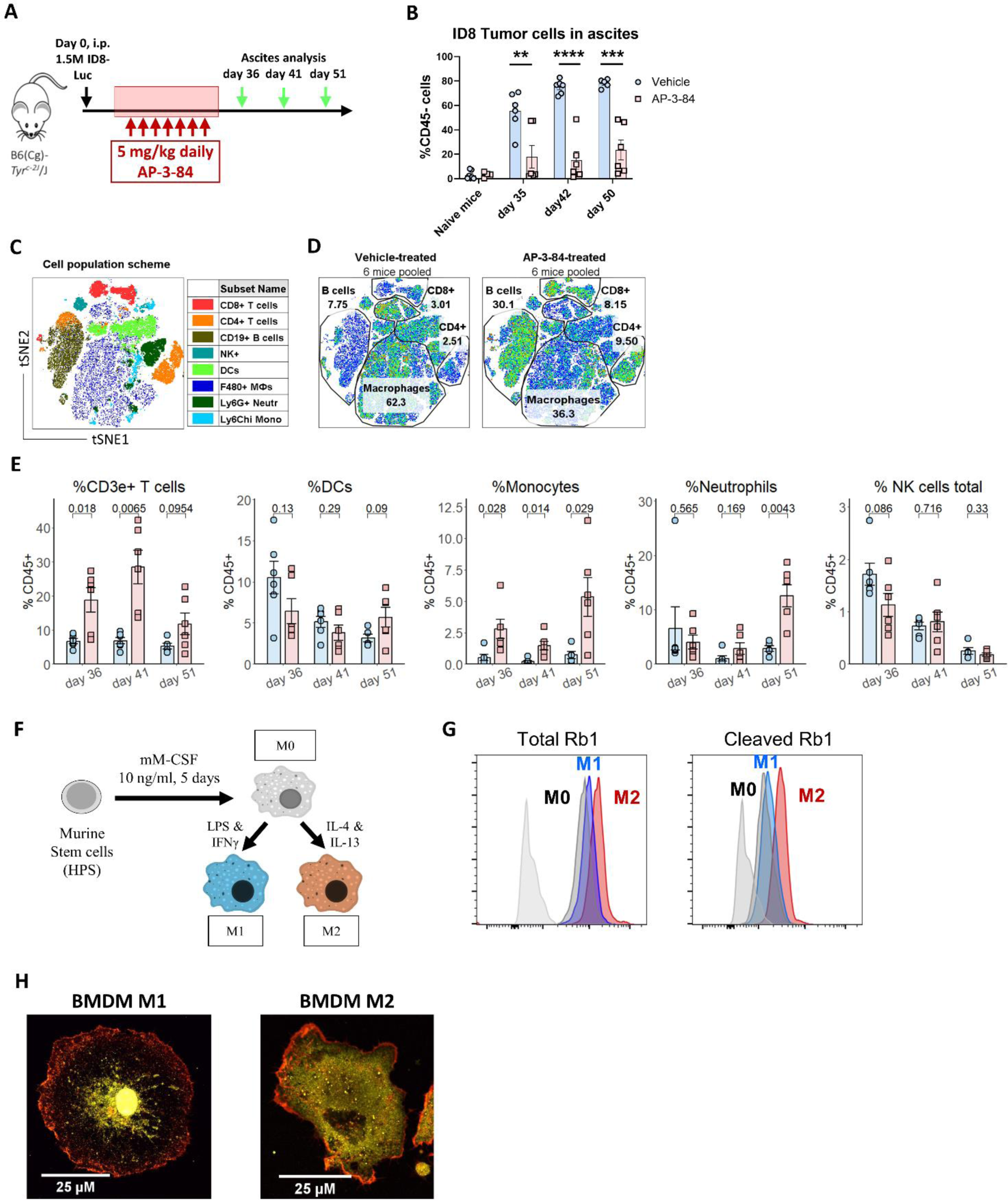
Therapeutic myeloid Rb targeting changes the composition of immune cell populations in the ascites of ID8-bearing mice and preferentially affects M2 type polarized macrophages. **(A)** Experimental model of ID8 ovarian cancer. B6 albino mice were intraperitoneally injected with 1.5×10^6^ ID8-Luc ovarian cancer cells, treated with 7 daily injections of 5 mg/kg AP-3-84 starting on day 27 with the following weekly tumor burden measurement by IVIS and ascites analysis by flow cytometry soon after the treatment end on day 36, then on day 41 and at progressed disease stage on day 51. **(B)** Frequency of tumor CD45– cells in total ascites of ID8-bearing mice at different time-points measured by flow cytometry. Peritoneal washes from intact naïve mice were used as a negative control. n=4-7 per time-point per group. **(C,D)** tSNE analysis was conducted on ascites CD45+ cells collected after AP-3-84 treatment on day 36. Schematic distribution (**C**) and the comparison of frequencies of major leukocyte populations in ascites of vehicle vs AP-3-84-treated mice (**D**) are shown. n=6 per condition. **(E)** On the indicated days tumor ascites was collected from AP-3-84-treated and control mice and flow cytometry quantification of the specified immune cell populations was conducted, also shown in **Fig. 1D**. n=6 per condition. **(F)** Generation of Bone Marrow-Derived Macrophages (BMDMs). HPCs were isolated from mouse bone marrow by negative selection with magnetic beads and incubated in culture media with 10 ng/ml mM-CSF for 5 days to generate “M0” macrophages. After that cells were left untreated as a control M0 cells or polarized to M1 or M2 type with the addition of LPS+IFN-γ (M1) or IL-4+IL-13 (M2) cytokines respectively for 2 more days. **(G)** Flow cytometry comparison of total Rb and cleaved Rb expression in BMDMs polarized to M1 and M2 types. Representative experiment out of 3 independent experiments is shown. **(H)** Generated BMDMs of M1 and M2 type were analyzed by confocal microscopy for the expression and distribution of Rb1 (yellow) and actin (red). Means +/– SEM are shown. Statistical significance was assessed by unpaired Student t-test; corresponding p-values are shown as exact numbers or indicated as *p<0.05, **p<0.01, ***p<0.001, ****p<0.0001.

**Supplementary Fig. S4.**
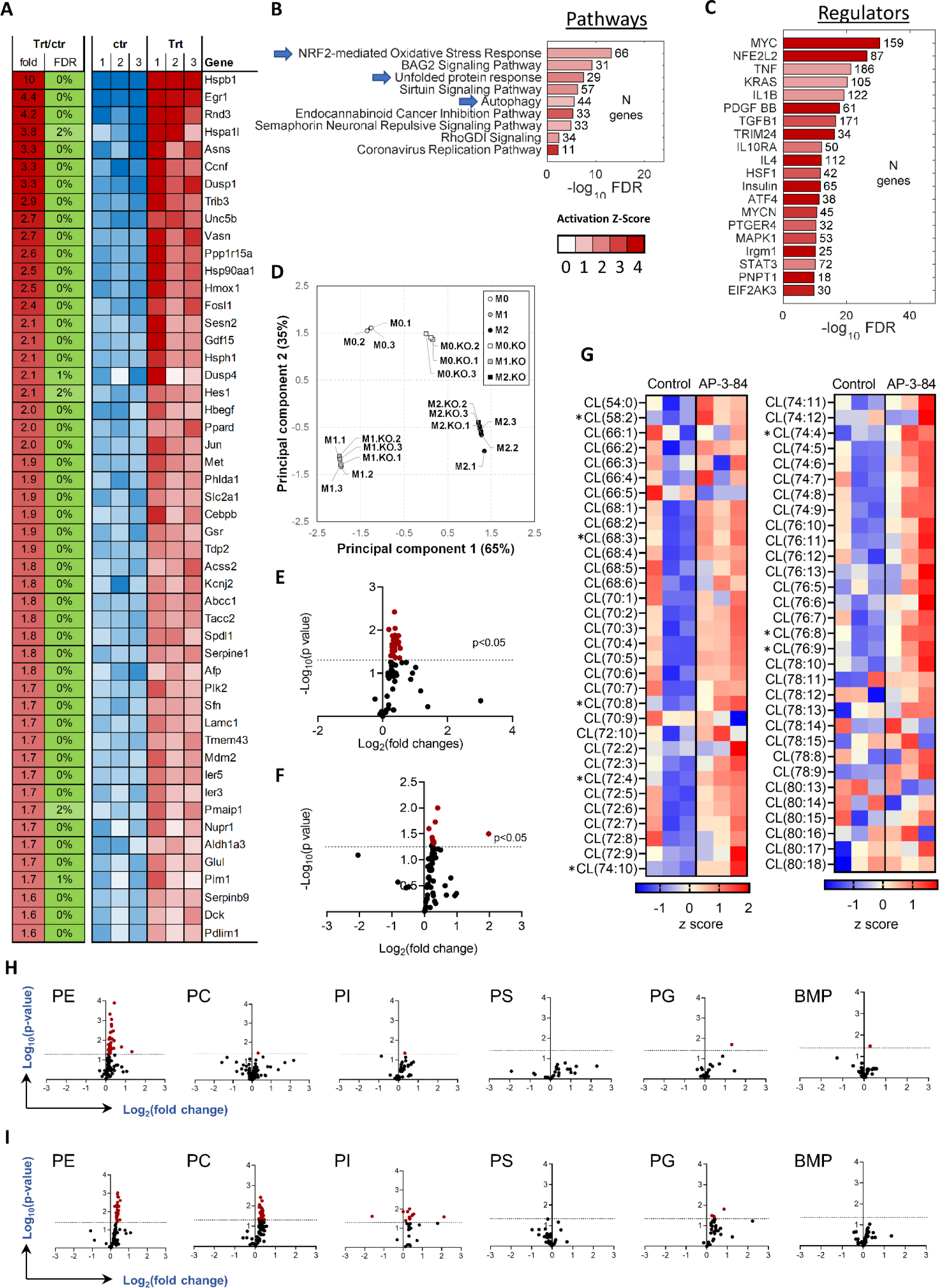
Myeloid Rb targeting initiates dramatic changes in macrophage gene expression profile and lipid composition inducing the major stress response pathways. Ascites macrophages from ID8 tumor-bearing mice (day 42 after ID8 tumor cell injection), thioglycolate macrophages or BMDM from wild type or p53 KO mice polarized to M1 or M2 type were generated and isolated, then treated *in vitro* with 10 uM AP-3-84 Rb modulator for 4 hours, washed, lysed and their gene expression was analyzed by RNA sequencing (n=3 per condition). **(A)** Top 50 genes in thioglycolate-elicited macrophages induced by AP-3-84 treatment are shown (false discovery rate, FDR<5%). **(B,C)** Ingenuity Pathways Analysis (IPA analysis) of RNA-Seq data for thioglycolate macrophages treated with AP-3-84. Canonical pathways **(B)** and upstream regulators (**C**) that were most dramatically activated upon AP-3-84 treatment are shown. Blue arrows highlight results of interest. **(D)** p53 KO BMDMs polarized to M0, M1 or M2 type (in triplicates indicated by “KO.1”, “KO.2”, and “KO.3”, respectively) were prepared, processed and projected on the same principal component analysis as in **Fig. 4A**. **(E)** Cardiolipin species analysis by LC-MS/MS in thioglycolate-elicited wild-type F4/80+ macrophages upon AP-3-84 treatment. Volcano plot demonstrating the differences in the contents of individual cardiolipin species is shown (n=3 per group). **(F,G)** ID8 ascites F4/80+ macrophages were isolated from ID8-bearing mice and incubated with 10 uM AP-3-84 *in vitro* for 4 hours, lysed and cardiolipin species were analyzed by LC-MS. Volcano plot (**F**) and heatmap (**E**) demonstrating the differences in the contents of individual molecular species of cardiolipins are shown. The horizontal dotted-line in (**F**) indicates p = 0.05 with points above the line having p < 0.05 and points below the line having p > 0.05. Data in (**E**) is presented as heat map auto-scaled to z scores and coded blue (low values) to red (high values). Asterisks show the species were significantly changed (p<0.05, t-test) after treatment with AP-3-38. n=3 per condition. **(H,I)** ID8 ascites F4/80+ macrophages **(H)** and thioglycollate-induced F4/80+ macrophages (**I**) were isolated and incubated with 10 uM AP-3-84 *in vitro* for 4 hours, lysed and lipid species of different classes were analyzed by LC-MS. Volcano plots demonstrating the differences in the contents of individual molecular lipid species are shown across major classes of phospholipids: PE, phosphatidylethanolamine, PC, phosphatidylcholine, PI, phosphatidylinositol, PS, phosphatidylserine, PG, phosphatidylglycerol, BMP, bis-monoacylglycerophosphate. The horizontal dotted line indicates p = 0.05 with points above the line having p < 0.05 and points below the line having p > 0.05. n=3 per condition. Summarized statistics for the corresponding populations are shown *p<0.05, **p<0.01, ***p< 0.001, **** p<0.0001 by unpaired Student t-test comparison.

**Supplementary Fig. S5.**
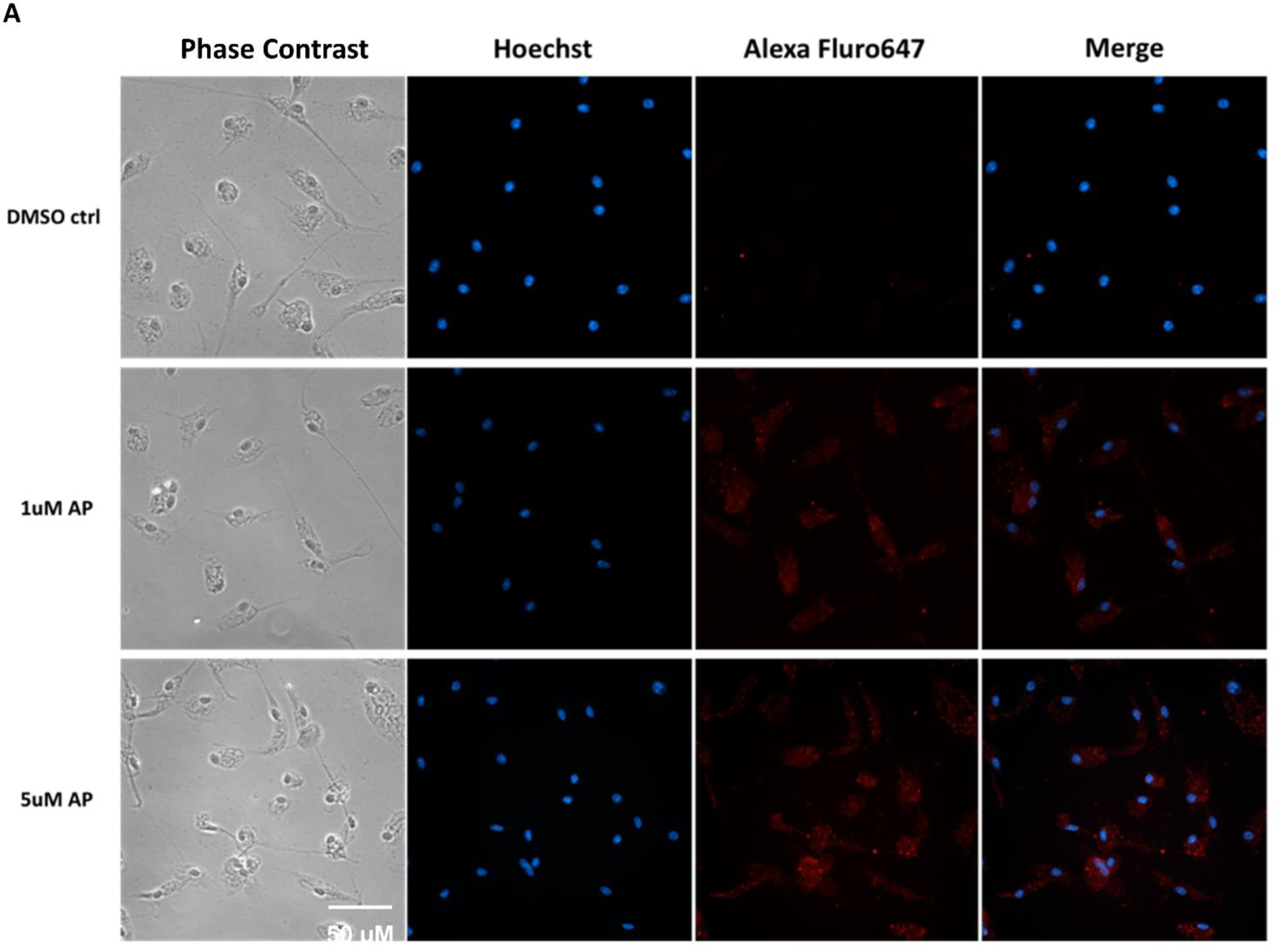
Therapeutic Rb targeting upregulates p53 expression in macrophages. (**A**) Thioglycollate-induced macrophages were seeded onto the wells of chamber slides for culture on day 1. On day 2, cells were treated with DMSO, 1 µM or 5 µM AP-3-84 overnight with the following immunofluorescence analysis. Representative data from at least three independent experiments is shown.

**Supplementary Fig. S6.**
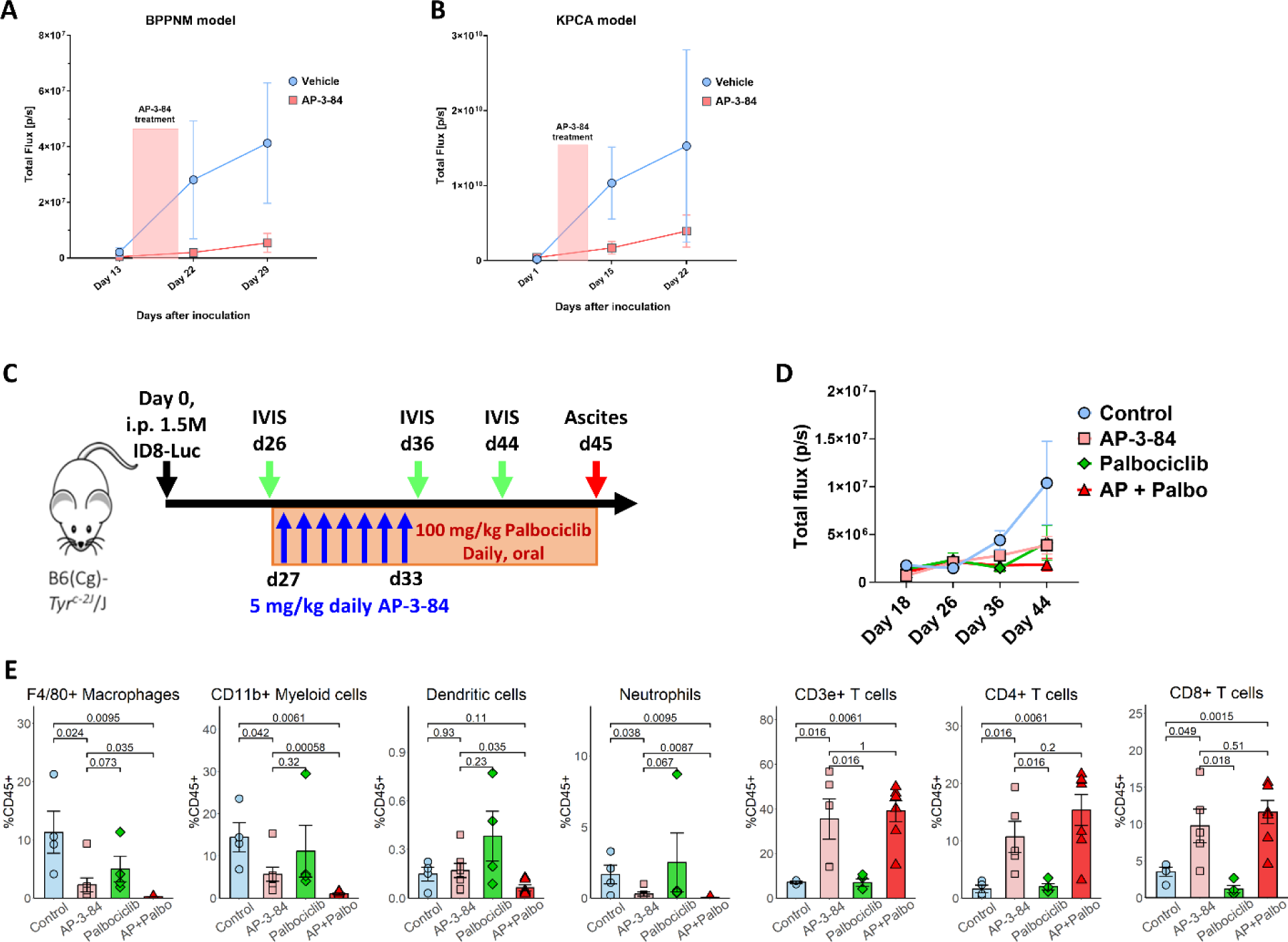
Therapeutic Rb targeting delays tumor growth in various ovarian cancer models and demonstrates the effects non-redundant to CDK4/6 palbociclib inhibitor. (**A,B**) BPPNM or KPCA ovarian cancer cells were injected intraperitoneally in mice with the following 7-day AP-3-84 treatment. Tumor burden was measured by IVIS on indicated days. n=5-8 per group. (**C-E**) ID8 ovarian cancer model was established and treated with AP-3-84 as shown in (**C**), similar to the setting in **Supplementary** Fig. 3A. The additional group was treated with palbociclib alone or in combination with AP-3-84 daily starting from day 27 until the end of the experiment. Tumor growth was monitored by IVIS measurements (**D**). On day 45 mice were sacrificed, their tumor ascites were collected and analyzed for immune cell composition by flow cytometry (**E**). n=4-6 per group.

**Supplementary Fig. S7.**
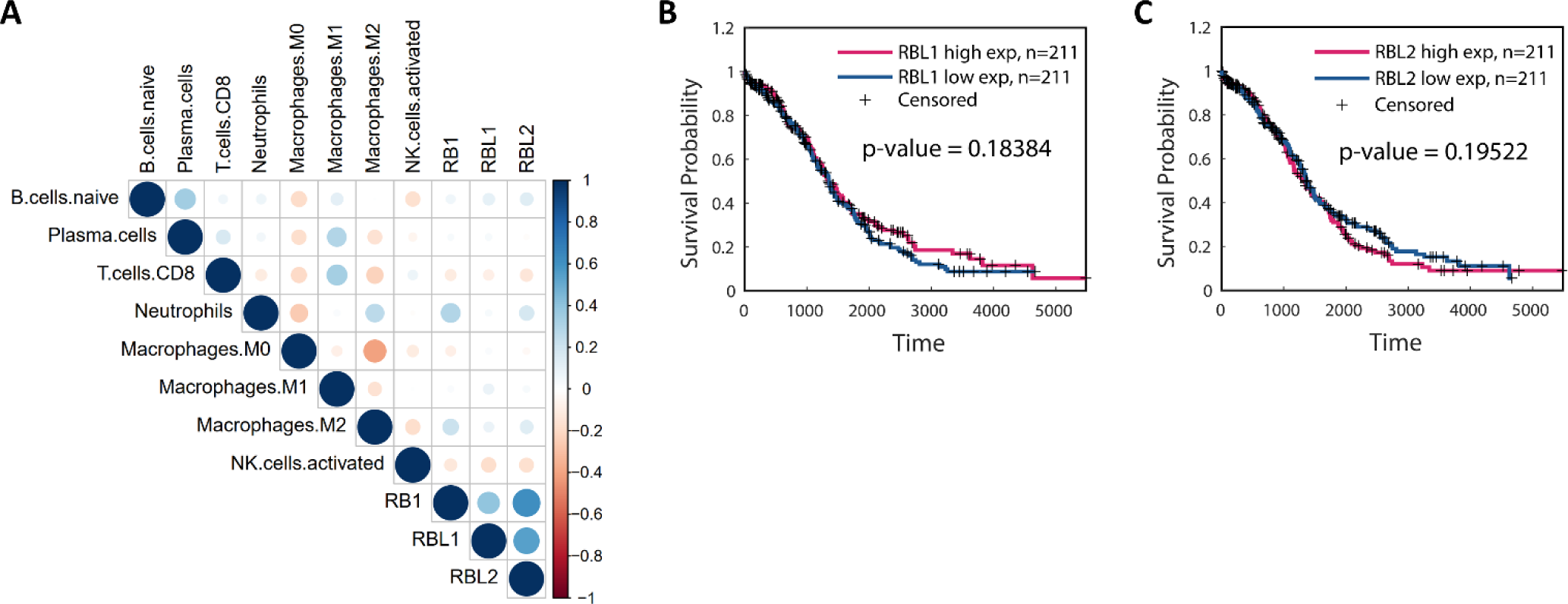
Comparison of patient survival outcome with different expression levels of RBL1/RBL2 in tumor. (**A**) Correlation heatmap showing the rank correlation coefficient when correlating RB1, RBL1 and RBL2 to CIBERSORT-inferred immune cell abundances through TCGA ovarian cancer samples. (**B,C**) Kaplan Meier survival curves comparing between TCGA ovarian cancer patients with higher and lower expression of RBL1 **(B)** or RBL2 (**C**). Low and high levels are separated by the respective median values. The log-rank p-values are indicated.

## Supplemental Tables

**Supplementary Table S1. The effect of HDAC inhibitors on Rb-HDAC1 interaction.**

Titrations of HDAC1 inhibitors were added using acoustic liquid handler by direct dilution such that the final amount of DMSO was 100 nL in each well. Final concentrations ranged from 0.32 to 10000 nM. 5 nM GST-Rb 380-785 pre-incubated with anti-GST-Tb HTRF donor antibody in assay buffer was added to each well. After a 15 min pre-incubation, 5 nM HDAC1-FLAG pre-incubated with anti-FLAG-Alexa488 HTRF acceptor antibody in assay buffer was added to each well. After a 1 hr incubation, the HTRF signal at 520 nm and 620 nm was measured using the ClarioStar plate reader.

**Supplementary Table S2. M0/M1/M2 signature genes.**

Gene expression in M0, M1 and M2-polarized BMDMs was analyzed by RNA Sequencing and most significantly different genes in each macrophage type shown in the **Supplementary Table S2** were used to generate M0/M1/M2 gene signature.

