## Supplementary Figures for "Targeting LxCxE cleft pocket of retinoblastoma protein in M2 macrophages inhibits ovarian cancer progression"

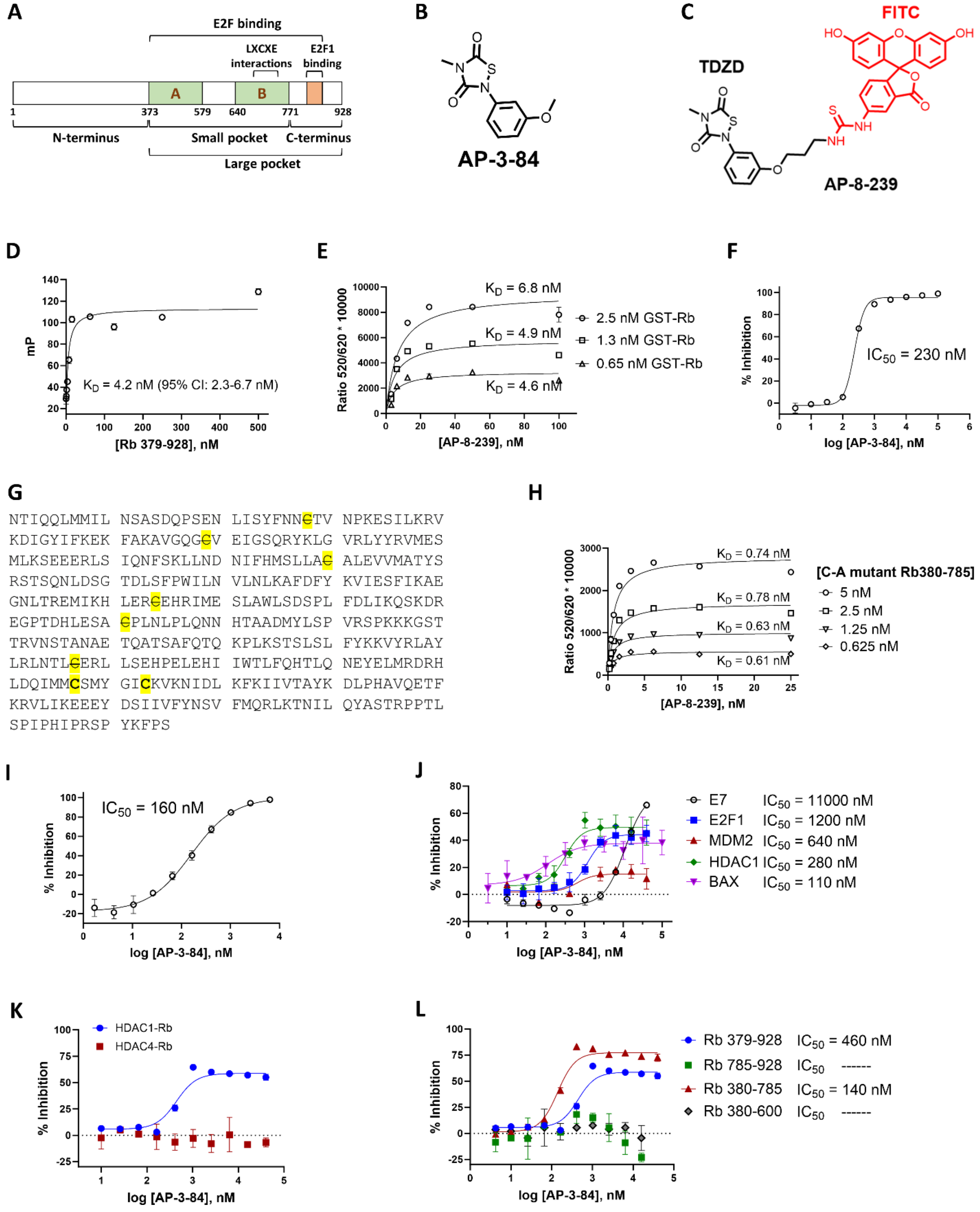

**Supplementary Fig. S1. Targeting the LxCxE cleft pocket of Rb protein with AP-3-84 modulates its interactions with the adaptor proteins.**

**(A)** Rb protein scheme according to (1).

**(B)** Chemical structure of AP-3-84.

**(C)** Chemical structure of AP-8-239 (AP-3-84-FITC) probe.

**(D)** AP-8-239 binds to untagged Rb (379-928), measured by fluorescence polarization (FP).

**(E)** AP-8-239 binds to GST-Rb (379-928) as measured by HTRF.

**(F)** Inhibition AP-8-239 binding to GST-Rb (379-928) by AP-3-84 compound. AP-3-84 was first pre-incubated with Rb, followed by AP-08-239 addition and HTRF measurement.

**(G)** Rb (380 – 785) protein sequence. All the highlighted cysteines were mutated to alanine (indicated with a strikethrough font) except Cys706 and Cys712.

**(H)** AP-8-239 binds to Cys706 and Cys712 of mutant Rb380-785, measured by HTRF as above.

**(I)** AP-3-84 inhibits AP-8-239 binding to C-A mutant 380-785 measured by HTRF as above.

**(J)** AP-3-84 inhibits the binding of Rb interacting proteins, measured by HTRF as above.

**(K)** AP-03-84 inhibits HDAC1, but not HDAC4 binding to mutant GST-Rb379-928.

**(L)** AP-3-84 inhibits HDAC1 binding to Rb by binding to the 380-785 region, measured by HTRF as above.

Mean +/–SD are shown (n=2-4 per condition). Representative experiment out of at least 3 repeats is shown.

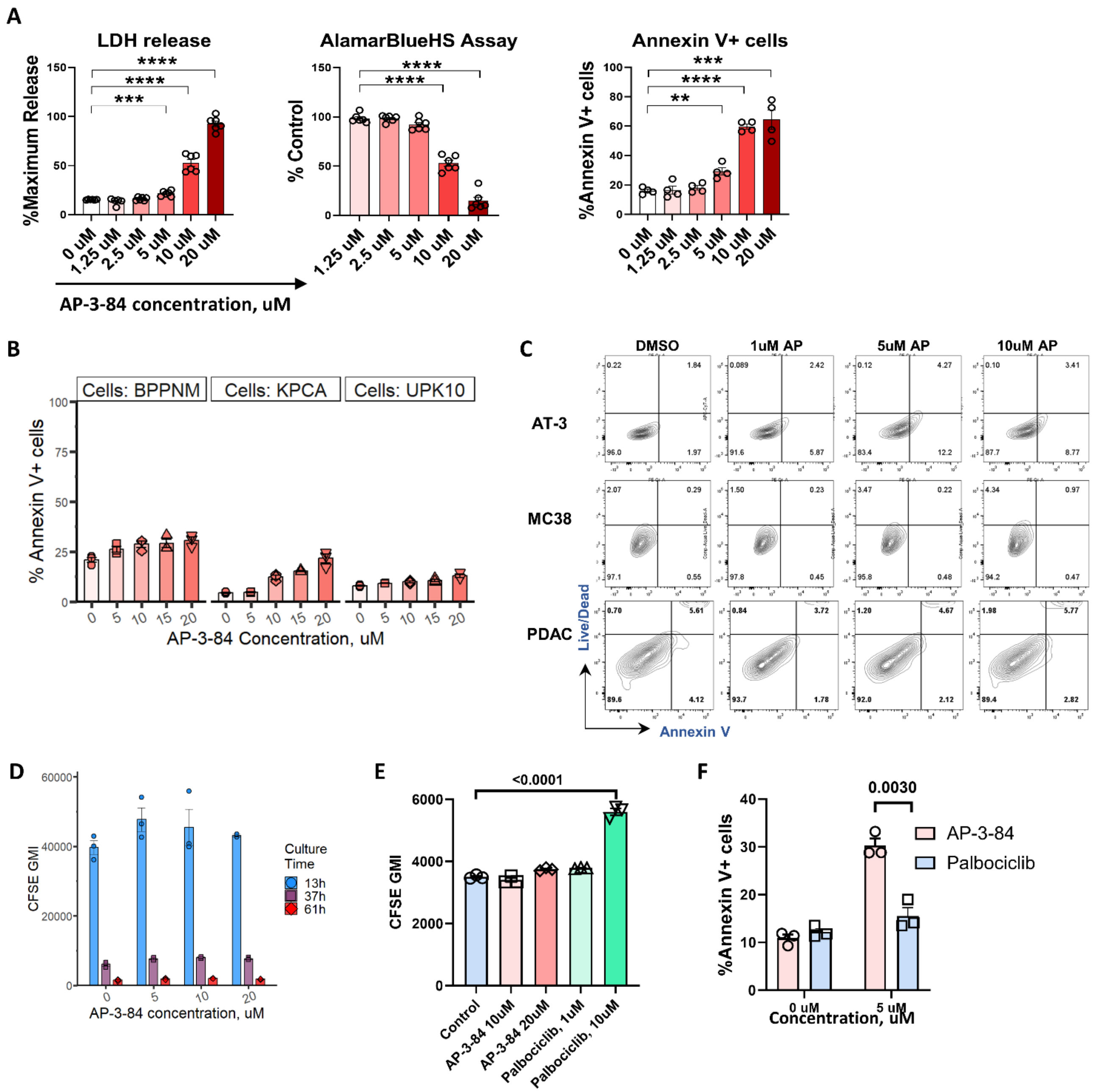


**Supplementary Fig. S2. Therapeutic myeloid Rb targeting preferentially affects macrophages, but not tumor cells and demonstrates the effects different from CDK4/6 palbociclib inhibitor.**

**(A)** Thioglycollate-induced macrophages were isolated with magnetic F4/80-beads from the peritoneal washes of mice 4 days after thioglycolate injection. Macrophages were incubated 16-

18 hours with different concentrations of AP-3-84. Viability of peritoneal macrophages was measured by LDH assay, AlamarBlueHS assay and Annexin V+/Live-Dead flow cytometry analysis. n=4-6 per condition.

**(B)** AP-3-84 effect on viability of different ovarian cancer cell lines: BPPNM, KPCA, UPK10. Cell treatment and annexin V staining were conducted as in **Fig. 1B**,**C**. n=3 per condition.

**(C)** AP-3-84 effect on viability of AT-3 mammary carcinoma, MC-38 colon cancer, and PDAC pancreatic cancer cells was tested as in (**C**) by Annexin V+/Live-Dead flow cytometry analysis.

**(D)** ID8 tumor cell proliferation in the presence of AP-3-84 at specified concentrations was measured by CFSE dilution assay after 13h, 37h and 61h of ID8 cell coincubation with AP-3-84. The similar drop of CFSE geometric mean intensity (CFSE GMI) during the cell culture is shown. n=3 per condition.

**(E)** The effect of 2-day treatment of ID8 tumor cells with the specified concentrations of AP-3-84 and palbociclib on cell proliferation measured by CFSE dilution assay. n=3 per condition.

**(F)** Ascites macrophages were isolated from ID8-bearing mice, incubated with the specified concentrations of AP-3-84 or palbociclib overnight and cell viability was measured by Annexin V/DAPI staining. n=3 per condition.

Means +/– SEM are shown with indicated exact p-values or **$p<0.01$, ***$p< 0.001$, **** $p<0.0001$ by unpaired Student t-test comparison.

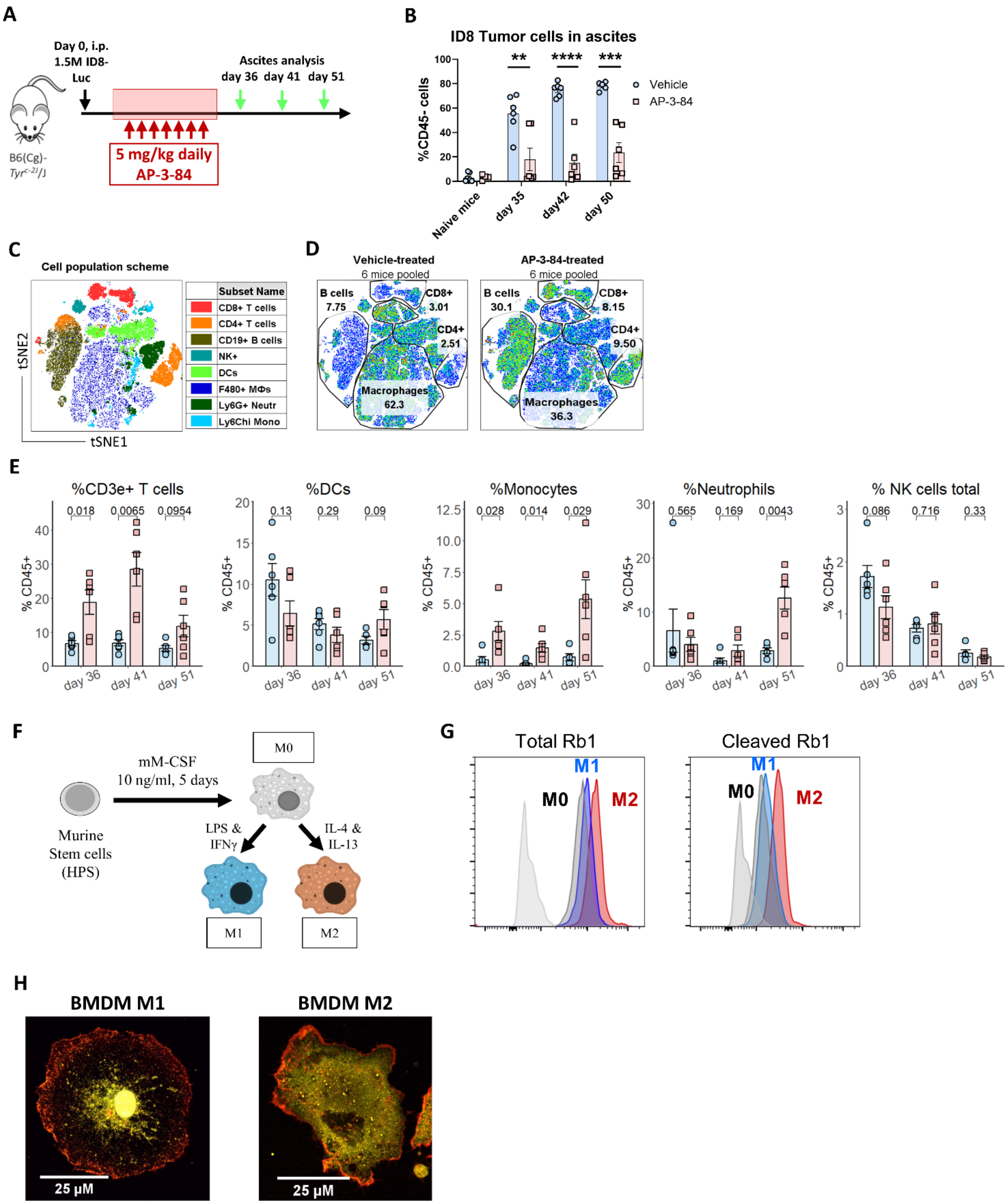

A
Day 0, i.p. 1.5M ID8-Luc
Ascites analysis
day 36 day 41 day 51
B6(Cg)-Tyr c-2J/J
5 mg/kg daily AP-3-84
B
ID8 Tumor cells in ascites
%CD45- cells
Vehicle
AP-3-84
Naive mice
day 35
day42
day 50
C
Cell population scheme
Subset Name
CD8+ T cells
CD4+ T cells
CD19+ B cells
NK+
DCs
F480+ MΦs
Ly6G+ Neutr
Ly6Chi Mono
tSNE2
tSNE1
D
Vehicle-treated
6 mice pooled
B cells 7.75
CD8+ 3.01
CD4+ 2.51
Macrophages 62.3
AP-3-84-treated
6 mice pooled
B cells 30.1
CD8+ 8.15
CD4+ 9.50
Macrophages 36.3
E
%CD3e+ T cells
0.018 0.0065 0.0954
%DCs
0.13 0.29 0.09
%Monocytes
0.028 0.014 0.029
%Neutrophils
0.565 0.169 0.0043
% NK cells total
0.086 0.716 0.33
% CD45+
day 36 day 41 day 51
F
mM-CSF
10 ng/ml, 5 days
M0
Murine Stem cells (HPS)
LPS & IFNγ
IL-4 & IL-13
M1
M2
G
Total Rb1
Cleaved Rb1
M0
M1
M2
H
BMDM M1
BMDM M2
25 μM
25 μM

Means +/– SEM are shown. Statistical significance was assessed by unpaired Student t-test; corresponding p-values are shown as exact numbers or indicated as *p<0.05, **p<0.01, ***p<0.001, ****p<0.0001.

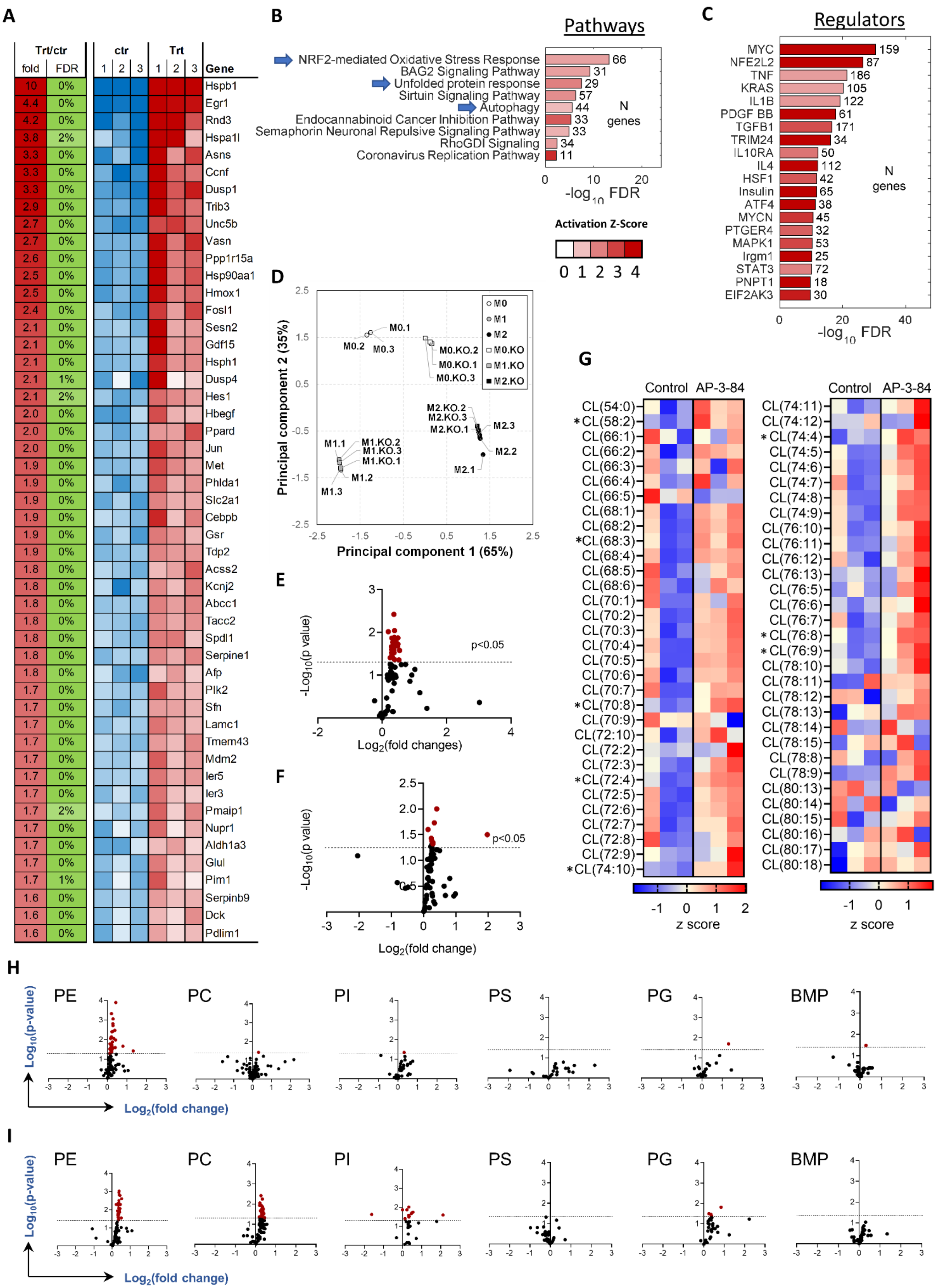

A
Trt/ctr
fold
FDR
ctr
Trt
1 2 3 1 2 3
Gene
10 0% Hspb1
4.4 0% Egr1
4.2 0% Rnd3
3.8 2% Hspa1l
3.3 0% Asns
3.3 0% Ccnf
3.3 0% Dusp1
2.9 0% Trib3
2.7 0% Unc5b
2.7 0% Vasn
2.6 0% Ppp1r15a
2.5 0% Hsp90aa1
2.5 0% Hmox1
2.4 0% Fosl1
2.1 0% Sesn2
2.1 0% Gdf15
2.1 0% Hsph1
2.1 1% Dusp4
2.1 2% Hes1
2.0 0% Hbegf
2.0 0% Ppard
2.0 0% Jun
1.9 0% Met
1.9 0% Phlda1
1.9 0% Slc2a1
1.9 0% Cebpb
1.9 0% Gsr
1.9 0% Tdp2
1.8 0% Acss2
1.8 0% Kcnj2
1.8 0% Abcc1
1.8 0% Tacc2
1.8 0% Spdl1
1.8 0% Serpine1
1.8 0% Afp
1.7 0% Plk2
1.7 0% Sfn
1.7 0% Lamc1
1.7 0% Tmem43
1.7 0% Mdm2
1.7 0% Ier5
1.7 0% Ier3
1.7 2% Pmaip1
1.7 0% Nupr1
1.7 0% Aldh1a3
1.7 0% Glul
1.7 1% Pim1
1.6 0% Serpinb9
1.6 0% Dck
1.6 0% Pdlim1
B
Pathways
NRF2-mediated Oxidative Stress Response 66
BAG2 Signaling Pathway 31
Unfolded protein response 29
Sirtuin Signaling Pathway 57
Autophagy 44
Endocannabinoid Cancer Inhibition Pathway 33
Semaphorin Neuronal Repulsive Signaling Pathway 33
RhoGDI Signaling 34
Coronavirus Replication Pathway 11
N genes
0 10 20
-log10 FDR
Activation Z-Score
0 1 2 3 4
C
Regulators
MYC 159
NFE2L2 87
TNF 186
KRAS 105
IL1B 122
PDGF BB 61
TGFB1 171
TRIM24 34
IL10RA 50
IL4 112
HSF1 42
Insulin 65
ATF4 38
MYCN 45
PTGER4 32
MAPK1 53
Irgm1 25
STAT3 72
PNPT1 18
EIF2AK3 30
N genes
0 20 40
-log10 FDR
D
Principal component 2 (35%)
Principal component 1 (65%)
M0
M1
M2
M0.KO
M1.KO
M2.KO
M0.1
M0.2
M0.3
M0.KO.2
M0.KO.1
M0.KO.3
M2.KO.2
M2.KO.3
M2.KO.1
M2.3
M2.2
M2.1
M1.1
M1.KO.2
M1.KO.3
M1.KO.1
M1.2
M1.3
E
-Log10(p value)
Log2(fold changes)
p<0.05
F
-Log10(p value)
Log2(fold change)
p<0.05
G
Control AP-3-84
CL(54:0)
*CL(58:2)
CL(66:1)
CL(66:2)
CL(66:3)
CL(66:4)
CL(66:5)
CL(68:1)
CL(68:2)
*CL(68:3)
CL(68:4)
CL(68:5)
CL(68:6)
CL(70:1)
CL(70:2)
CL(70:3)
CL(70:4)
CL(70:5)
CL(70:6)
CL(70:7)
*CL(70:8)
CL(70:9)
CL(72:10)
CL(72:2)
CL(72:3)
*CL(72:4)
CL(72:5)
CL(72:6)
CL(72:7)
CL(72:8)
CL(72:9)
*CL(74:10)
z score
Control AP-3-84
CL(74:11)
CL(74:12)
*CL(74:4)
CL(74:5)
CL(74:6)
CL(74:7)
CL(74:8)
CL(74:9)
CL(76:10)
CL(76:11)
CL(76:12)
CL(76:13)
CL(76:5)
CL(76:6)
CL(76:7)
*CL(76:8)
*CL(76:9)
CL(78:10)
CL(78:11)
CL(78:12)
CL(78:13)
CL(78:14)
CL(78:15)
CL(78:8)
CL(78:9)
CL(80:13)
CL(80:14)
CL(80:15)
CL(80:16)
CL(80:17)
CL(80:18)
z score
H
PE PC PI PS PG BMP
Log10(p-value)
Log2(fold change)
I
PE PC PI PS PG BMP
Log10(p-value)
Log2(fold change)

Summarized statistics for the corresponding populations are shown *$p<0.05$, **$p<0.01$, ***$p< 0.001$, **** $p<0.0001$ by unpaired Student t-test comparison.

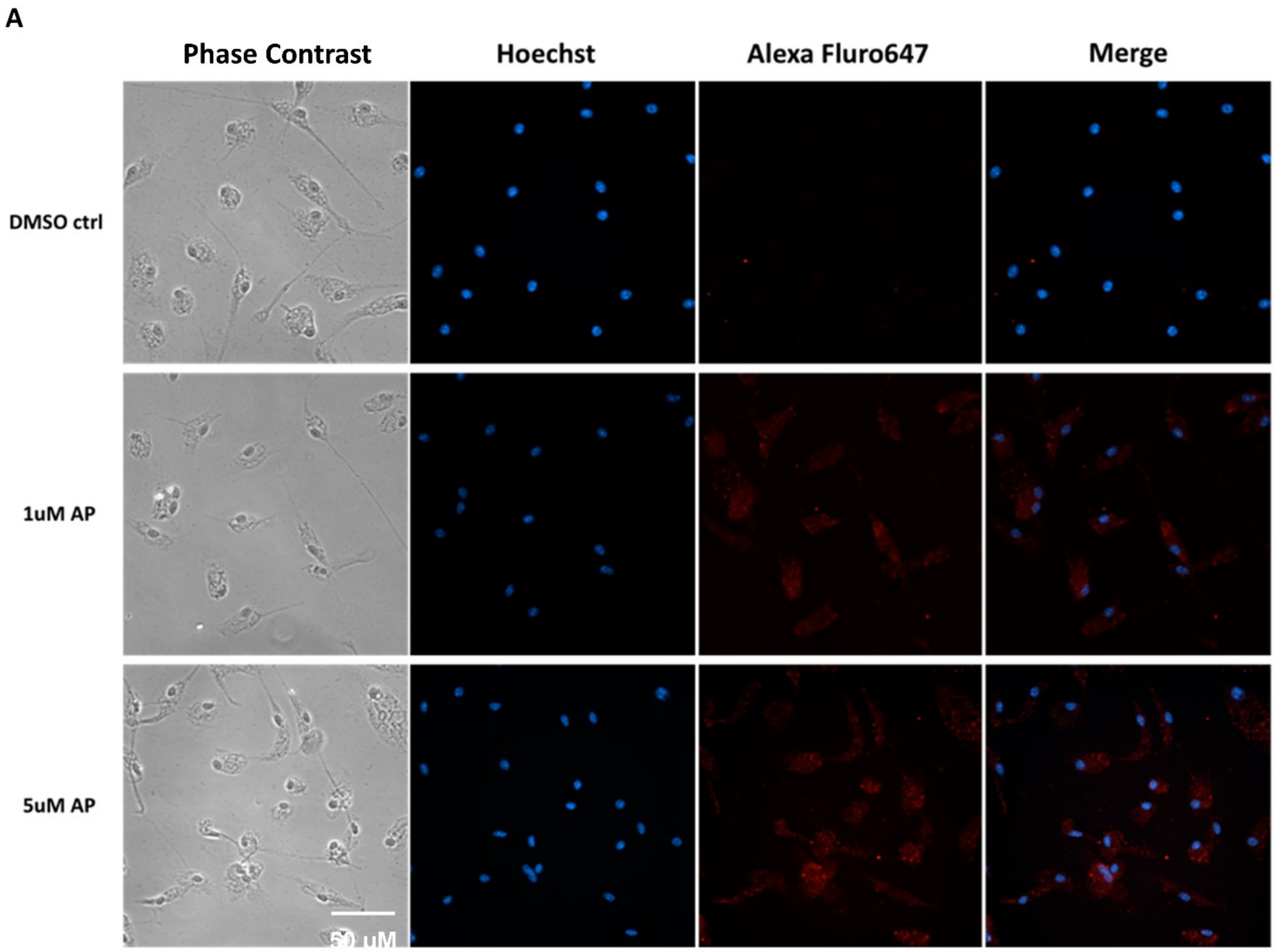


**Supplementary Fig. S5. Therapeutic Rb targeting upregulates p53 expression in macrophages.**

(**A**) Thioglycollate-induced macrophages were seeded onto the wells of chamber slides for culture on day 1. On day 2, cells were treated with DMSO, 1 μM or 5 μM AP-3-84 overnight with the following immunofluorescence analysis. Representative data from at least three independent experiments is shown.

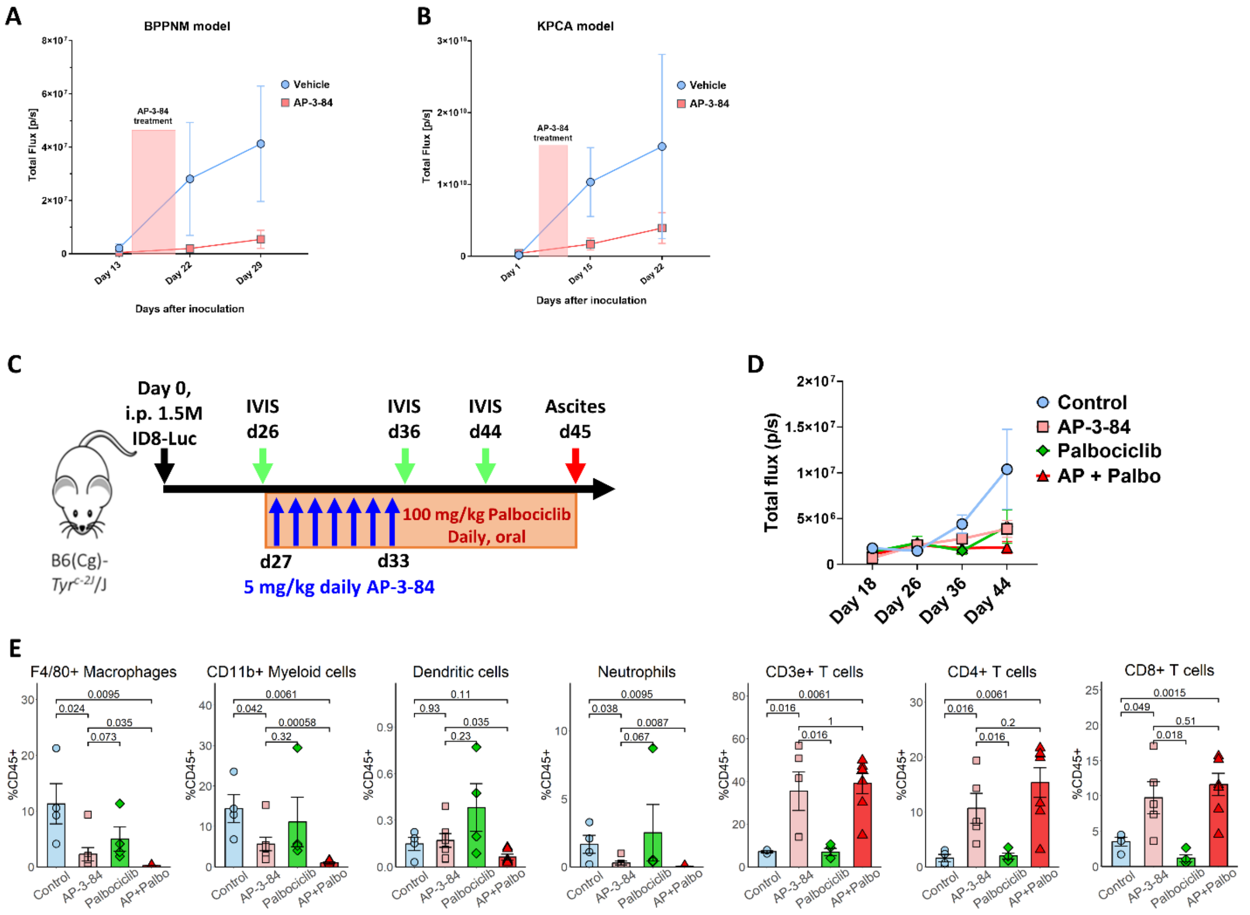


**Supplementary Fig. S6. Therapeutic Rb targeting delays tumor growth in various ovarian cancer models and demonstrates the effects non-redundant to CDK4/6 palbociclib inhibitor.** (**A,B**) BPPNM or KPCA ovarian cancer cells were injected intraperitoneally in mice with the following 7-day AP-3-84 treatment. Tumor burden was measured by IVIS on indicated days. n=5-8 per group.

(**C-E**) ID8 ovarian cancer model was established and treated with AP-3-84 as shown in (**C**), similar to the setting in **Supplementary Fig. 3A**. The additional group was treated with palbociclib alone or in combination with AP-3-84 daily starting from day 27 until the end of the experiment. Tumor growth was monitored by IVIS measurements (**D**). On day 45 mice were sacrificed, their tumor ascites were collected and analyzed for immune cell composition by flow cytometry (**E**). n=4-6 per group.

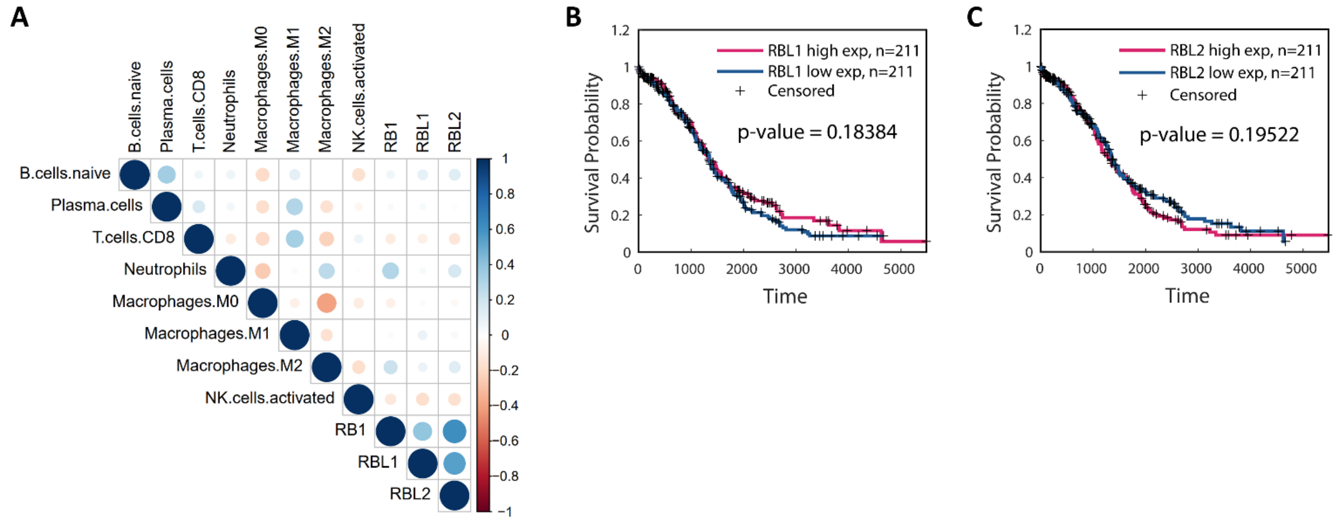


**Supplementary Fig. S7. Comparison of patient survival outcome with different expression levels of RBL1/RBL2 in tumor.**

(**A**) Correlation heatmap showing the rank correlation coefficient when correlating RB1, RBL1 and RBL2 to CIBERSORT-inferred immune cell abundances through TCGA ovarian cancer samples.

(**B,C**) Kaplan Meier survival curves comparing between TCGA ovarian cancer patients with higher and lower expression of RBL1 **(B)** or RBL2 (**C**). Low and high levels are separated by the respective median values. The log-rank p-values are indicated.

### Supplemental Tables

**Supplementary Table S1. The effect of HDAC inhibitors on Rb-HDAC1 interaction.**

Titrations of HDAC1 inhibitors were added using acoustic liquid handler by direct dilution such that the final amount of DMSO was 100 nL in each well. Final concentrations ranged from 0.32 to 10000 nM. 5 nM GST-Rb 380-785 pre-incubated with anti-GST-Tb HTRF donor antibody in assay buffer was added to each well. After a 15 min pre-incubation, 5 nM HDAC1-FLAG pre-incubated with anti-FLAG-Alexa488 HTRF acceptor antibody in assay buffer was added to each well. After a 1 hr incubation, the HTRF signal at 520 nm and 620 nm was measured using the ClarioStar plate reader.

**Supplementary Table S2. M0/M1/M2 signature genes.**

Gene expression in M0, M1 and M2-polarized BMDMs was analyzed by RNA Sequencing and most significantly different genes in each macrophage type shown in the **Supplementary Table S2** were used to generate M0/M1/M2 gene signature.
